# p53 mediated regulation of LINE1 retrotransposition derived R-loops

**DOI:** 10.1101/2024.04.12.589154

**Authors:** Pratyashaa Paul, Arun Kumar, Astik Kumar De, Ankita Subhadarsani Parida, Gauri Bhadke, Satyajeet Khatua, Fizalin Pattanayak, Bhavana Tiwari

**Affiliations:** Department of Biological Sciences, Indian Institute of Science Education and Research Berhampur, India

**Keywords:** Keywords, p53, L1/LINE1, R-loops, RNA-DNA hybrids, Cis R-loop, Trans R-loop, Transposons, HDAC inhibitors

## Abstract

Long Interspersed Nuclear Element 1 (LINE1/L1) retrotransposons, comprising around 17% of the human genome, typically remain quiescent in healthy somatic cells but become activated in various cancer types. Our recent investigation reveals that p53 silences L1 transposons in human somatic cells, potentially constituting a tumor suppressive pathway. In this study, we demonstrate that p53 silences both L1mRNA-gDNA (cis L1 R-loops) and L1mRNA-cDNA hybrids (trans L1 R-loops) formed during retrotransposition. The activation of L1 transposons by HDAC inhibitors (HDACi) led to accumulation of these cis and trans L1 R-loops in p53^-/-^ cells, which were mitigated by treatment with a reverse transcriptase inhibitor. Furthermore, p53 established re-silencing of hyperactivated L1 transposons induced by HDACi. The p53-mediated restoration of silencing was accompanied by recruiting histone repressive marks specifically H3K9me3 and H3K27me3 and inhibiting the deposition of H3K4me3 and H3K9ac marks at the L1 promoter. This study elucidates a novel role of p53 in regulating the formation of RNA-DNA hybrids, a pivotal intermediate component of retrotransposition, and initiating the suppression of hyperactivated L1 elements. These findings underscore the significance of p53 in preserving genome stability through the regulation of L1-derived R-loops.

**In Brief:** The role of L1 transposon derived L1mRNA-cDNA hybrids; an intermediate product formed during retrotransposition, in DNA damage and inflammation is not clear. Paul et al. reveals that p53 prevents L1cDNA derived RNA-DNA hybrids to control DNA damage and activation of inflammatory genes. The findings also elucidate the role of p53 in initiating the repression of hyperactivated transposons by facilitating the recruitment of epigenetic repressive marks and preventing the deposition of activating marks at L1-5’UTR.

**Highlights:**

1. p53 loss facilitates accumulation of both cis (L1mRNA-gDNA) and trans (L1mRNA-cDNA) forms of L1 R-loops.
2. The youngest, actively retrotransposing full-length L1s contribute to the formation of trans (L1mRNA-cDNA) R-loops.
3. p53 aids immediate L1 re-silencing by restoring deposition of epigenetic repressive and inhibition of activating marks.
4. Reverse transcriptase inhibitor prevents L1 mediated DNA damage.

**Subject Categories**: L1/LINE1, p53, Retrotransposons, RNA-DNA hybrids, Cis R loops, Trans R-loop, L1/LINE1

**Graphical:** 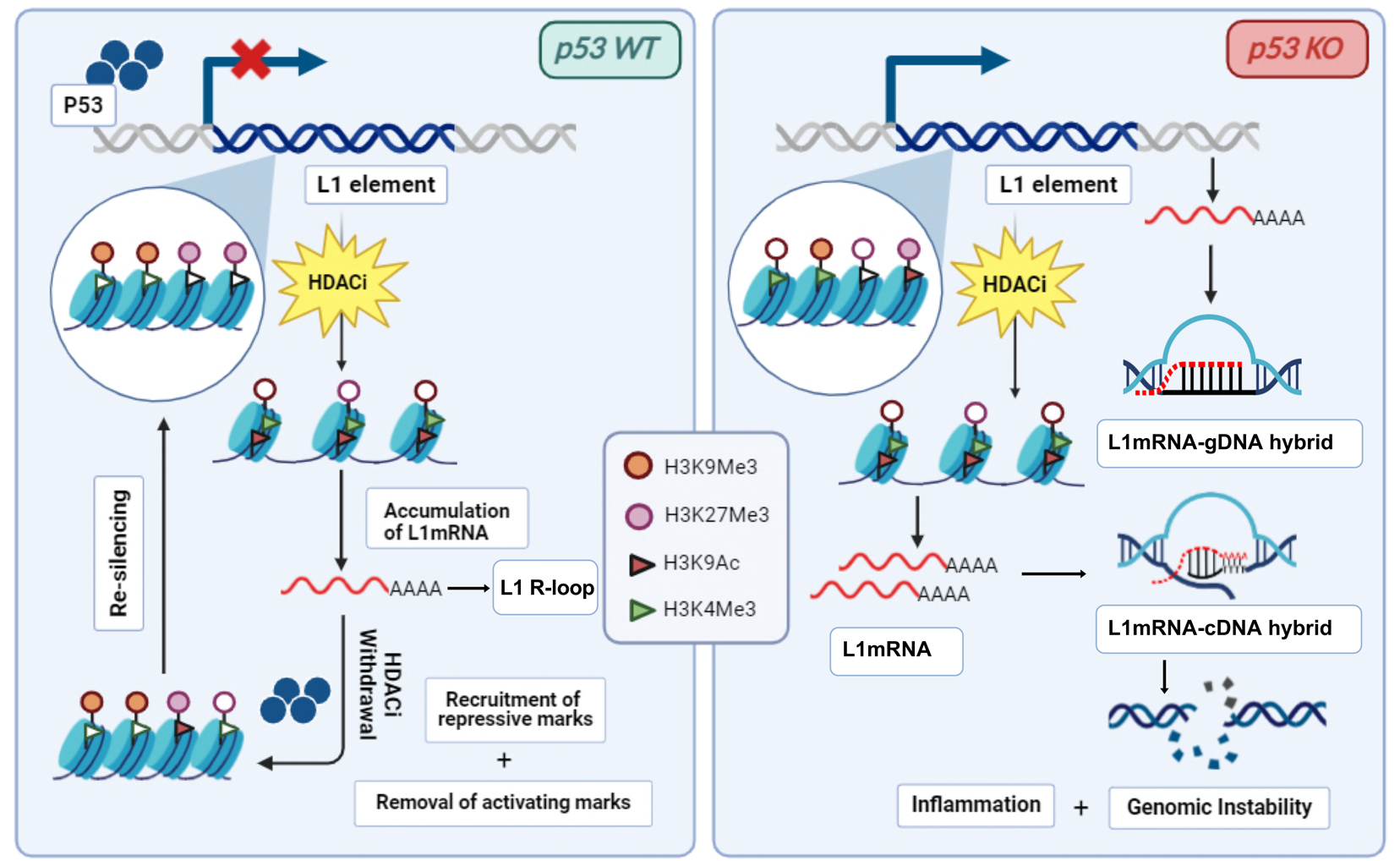

## Introduction

p53 is the most commonly mutated tumour suppressor gene ^1^ and activates its downstream effectors to regulate cell cycle, apoptosis and senescence pathways to maintain genome integrity ^2^. Recent studies have shown that p53 represses mobile genetic elements, suggesting a new axis of tumor suppression ^3–6^. p53 mediated retroelement repression is conserved across various species, including flies, fish, and human somatic cell lines ^7–10^. Employing CRISPR-mediated wild-type (wt) and p53 knockout (p53^-/-^) cells, we have identified p53 binding motifs within the 5’UTR of L1 retrotransposons, and work from others predicted in Alu sequences ^11–13^. By utilizing the synthetic L1-5’UTR-eGFP reporter, we demonstrated that p53 maintains repression of L1 within human somatic cell lines by facilitating the deposition of epigenetic repressive marks at the 5’UTR promoter to exert its repressive function ^9^. Several other studies have underscored a direct correlation between increased retroelement expression and p53 mutations within cancer contexts ^5,14,15^. These collective observations prompted an exploration into the potential involvement of p53 in initiating the silencing of activated L1 transposons and delineating the underlying mechanisms of *de novo* silencing. To address this, we employed HDAC inhibitors Trichostatin (TSA) and Sodium Butyrate (NaB) drugs that are known to induce repetitive elements, including L1 transposons on wt and p53^-/-^ cells ^16–18^. We examined the role of p53 in L1 resilencing function, a process that mirrors the epigenetic reprogramming observed in early embryos and primordial germ cells *in vivo*.

The initiation of L1 retrotransposition process involves transcription followed by the translation of two essential L1-encoded open reading frame proteins (ORFps); ORF1p and ORF2p. The combined efforts of ORF1p and ORF2p facilitate L1 retrotransposition and are linked to various diseases ^19–23^. These proteins are crucial for the reverse transcription of L1cDNA and its subsequent integration. During mobilization, these ORFs are transcribed by RNA pol II, leading to the formation of a L1mRNA-gDNA hybrid as a transcriptional intermediate, if these structures persists can lead to the formation of ‘cis R-loops’. R-loops represent three-stranded nucleic acid structures composed of an RNA/DNA hybrid alongside a displaced single-stranded DNA (ssDNA) and these can lead to either physiological or pathological outcomes. L1 encoded ORF2 protein contains two functional domains essential for target-primed reverse transcription: the Endonuclease (EN) domain creates a single-stranded DNA nick for integration, and the Reverse Transcriptase (RT) domain primes complementary DNA synthesis based on the L1mRNA template. The recent structural findings on ORF2p identified domains that show extensive interactions with RNA templates ^24^. The reverse transcription function of ORF2p generates a L1mRNA-cDNA hybrid structure, hence referred to as ‘trans R-loop’ intermediate structure comprising a free and un-nicked DNA segment, a nicked DNA, the newly synthesized cDNA, and L1mRNA serving as the template for reverse transcription, probably the four stranded non-canonical form of R-loops. L1 derived cDNA acts as triggers for type 1 interferon response, leading to the formation of inflammasomes ^25^. Ectopic accumulation of (cis) R loops, particularly those arising from transcriptional events, is associated with various genomic and epigenetic instabilities ^26–28^. These R-loops are implicated in genome instability ^29^ and DNA damage through several mechanisms: initiating abnormal replication ^30^, inducing transcription stress, and facilitating collisions between transcription and replication processes ^26^. Consequently, the accumulation of cis and trans R-loops derived from L1 sequences can potentially lead to DNA damage and inflammation, an aspect of L1 pathogenesis that remains unexplored. L1-derived RNA/DNA hybrids may be exported to the cytosol via an as yet unidentified process, where they are detected by sensors such as endosomal TLR3 and the membrane-bound cGAS inflammatory pathway ^31^. The L1 retroelement has the capability to retrotranspose Alus ^32,33^ and SVAs ^34,35^ hence these elements can also participate in the generation of unscheduled R-loops in the genome. There have been reports of ∼100 L1-mediated insertions, the end product of retrotransposition, leading to genetic diseases. These retroelement insertions have been linked to the transmission of congenital disorders through the germ line, familial predispositions to cancer, as well as other conditions like β-thalassemias, and hemophilias ^36–38^. However, there is an increasing acknowledgment that active L1s in somatic tissues also present inherent oncogenic hazard ^3,5,15^.

Herein, we establish the role of p53 in controlling the intermediate components of L1, retrotransposition specifically L1mRNA-cDNA hybrids, referred to here as ’L1 trans R-loops’, formed during reverse transcription, and its role in genome instability. We present evidence indicating that the most recently evolved full-length retrotransposition-competent L1 elements are actively involved in the generation of trans R-loops. This assertion is supported by our analysis of DRIP-seq (DNA-RNA immunoprecipitation sequencing) data, along with validation conducted through DRIP-qCR (quantitative PCR) experiments. Furthermore, our findings reveal that the over-expression of the RT domain of L1-ORF2p led to a notable increase in trans R-loop accumulation, which was accompanied by heightened levels of DNA damage and inflammation. Interestingly, treatment with a reverse transcriptase inhibitor attenuated these elevations in DNA damage and inflammation markers, underscoring the oncogenic implications of L1-derived trans R-loops. We also demonstrate that p53 mediates *de novo* silencing of hyperactivated L1 retroelements employing HDAC inhibitor (HDACi) treated cancer cell lines. Using synthetic L1-5’UTR-eGFP wt and p53^-/-^ reporters, combining with HDACi exposure, we observed highly de repressed L1 transposons in both the genotypes. However, upon the withdrawal of HDACi drugs, the silencing of de repressed L1 transposons and its retrotransposition intermediates was immediate in wt p53 cells but not in p53^-/-^ cells. This restoration of L1 silencing in wt L1 reporter coincided with the recruitment of histone repressive marks and the removal of activating marks at the L1-5’UTR providing insights into potential therapeutic strategies for diseases associated with L1 activation.

## Results

### p53 controls L1 retrotransposition intermediary components

Recently, our research demonstrated that p53 plays a crucial role in suppressing L1 transposons at the transcription levels in human cell lines under non-stress conditions ^9^. Building upon these findings, our current study delves into the consequences of p53 deficiency on the regulation of intermediary by-products of L1 activity, specifically RNA-DNA hybrids formed during retrotransposition. Here, we noted heightened levels of L1 transcripts (**Figures** 1A and B) and ORF1 protein (**Figures** 1C and 1D) in both p53^-/-^ cell lines (A375 and U2OS) generated using CRISPR cas9 methodology **(Figures** S1A and B), consistent with our prior study ^4^. The immunofluorescence (IF) experiments in wt and p53^-/-^ cells with s9.6, a widely used antibody to visualize and detect the RNA-DNA hybrids of R-loops ^39^, showed an increased signal of cytoplasmic RNA-DNA hybrids in p53^-/-^ (**Figures** 1E and 1E’) as compared to the parental wt cells. Next, DNA-RNA immunoprecipitation (DRIP) assays were conducted utilizing the s9.6 antibody to investigate the propensity of L1 element in the participation of R-loops in p53^-/-^ cells (**Figure** 1F). In this experiment s9.6 antibody was used to precipitate RNA-DNA hybrids structures followed by quantification by qPCR using the primers that binds the DNA region of interest to measure the binding ^40^. Mapping experiments with DRIP-qPCR using primers spanning 5’UTR, ORF1, and ORF2 regions revealed that the L1-5’UTR region, which harbors RNA pol II promoter, was predominantly enriched in R-loops derived RNA-DNA hybrids, whereas the ORF2 region exhibited a modest enrichment. The specificity of our assay for detecting RNA-DNA hybrids was validated by pre-treating the samples with RNase H before the antibody pulldown (data not shown), resulting in the complete abolition of our ability to amplify the region. Given that the majority of the nuclear R-loop derived RNA-DNA hybrids was observed in the cytoplasm, we leveraged an alternative method to specifically probe R-loops. This involved the overexpression of RNaseH1 with D210N catalytic dead mutant (RNH-GFP) with the nuclear localization signal fused at the N-terminus in A375 wt and p53^-/-^ cells ^27,40,41^. Stable cell lines of RNH-GFP showed the similar GFP expression between these genotypes. qPCR experiments from GFP immunoprecipitated samples indicated stronger R-loop signal at L1-5’UTR and a weaker signal at the gene body L1-ORF2 in RNH-GFP cells, similar to s9.6 pull down data **(Figures 1G and 1H)**. Moreover, our analysis of DRIP-seq and Global Run-On sequencing (GRO-seq) data unveils a partial overlap of peaks between RNA-DNA hybrid sequences and nascent transcripts at the L1-5’UTR coordinates (cis R-loops). Conversely, no overlapping peaks were observed at the L1-ORF2 region in these two sequencing datasets (**Figure** 1I), suggesting that 5’UTR region demonstrates a higher tendency to form cis R-loops, likely due to nascent RNA annealing. Conversely, ORF2 region possibly be involved in generation of trans R-loops, originating from distant L1mRNAs.

**Figure 1.**
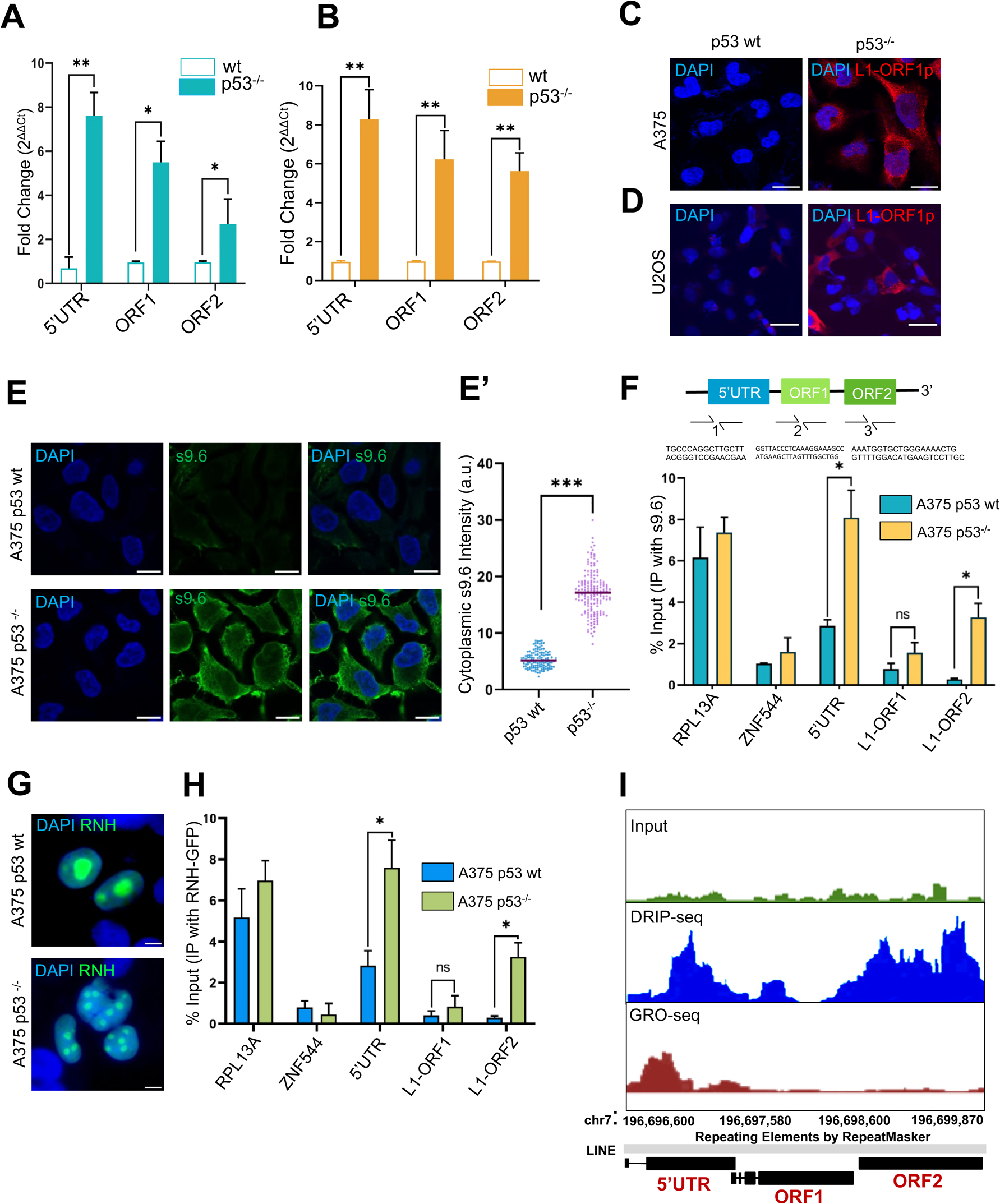
Elevation of L1 encoded intermediate components in p53^-/-^ cells: (A-B) qRT-PCR analyses was performed using the primers that detects consensus sequences of 5’UTR, L1ORF1 and L1ORF2 transcripts in wt and p53^-/-^ of (A) A375 and (B) U2OS cells. The transcript levels were normalized using ß-actin and relative fold change was calculated with respect to the parental wild type. The data shown are the averages of three biological replicates. *p<0.05, **p<0.01, (Mann-Whitney U test). (C-D) Representative confocal images display immunofluorescence staining for L1-ORF1p (Alexa Fluor 568, red) and DAPI (blue) in wt and p53^-/-^ cells of (C) A375 and (D) U2OS. Scale bars, 100 μm and 30 μm respectively. (E) Immunofluorescence images s9.6 stained (Alexa Fluor, 488) and DAPI (blue) A375 wt and p53^-/-^ cells. Scale bar,100 μm. Shown images are representative of three biological replicates. (E’) Quantification of s9.6 intensities of whole cell from A375 p53 wt and p53^-/-^ using fiji. Minimum 60 cells were analysed for RNA-DNA hybrid quantification from three biological replicates. *p<0.05, ns=non-significant (Mann-Whitney U test). (F) DRIP-qPCR was performed to measure R-loops in A375 p53 wt and p53^-/-^ cells. Immunoprecipitation carried out using the s9.6 antibody. Arrows at the schematic indicate to primers amplifying different regions of L1 sequence at 5’-UTR, ORF1 and ORF2). RPL13A was used a positive control for quantification of the R-loop locus and ZNF544 as negative locus. Enrichment of RNA-DNA hybrids in each region, normalized to input values. Data represent mean± S.D. from three biological experiments. *p<0.05, ns=non-significant (Mann-Whitney U test). (G) The representative fluorescent images show A375 wt and p53^-/-^ cells transfected with RNaseH1 D210N-GFP (RNH-GFP) construct. The nuclear foci indicate the expression of RNaseH1 D210N-GFP (RNH-GFP, green), nucleus (Blue). (H) DRIP-qPCR on RNaseH D210N-GFP stably transfected cells of A375 wt and p53^-/-^ cells using the primers detecting L1-5’UTR, ORF1 and ORF2 loci. RPL13A used as a positive control for the R-loop enrichment and ZNF544 as negative locus. Signal values of RNA-DNA hybrids immunoprecipitated in each region, normalized to input values. Data represent mean±S.D. from the three independent experiments. *p<0.05, ns=non-significant (Mann-Whitney U test). (I) This figure illustrates the comparison of DRIP-seq signals in the input control and immunoprecipitated sample, as well as the GRO-seq signal from the same cell line. The analyses from the dataset (PMID 26579211) focuses on a specific L1Hs genomic region. The green peak corresponds to the input of DRIP-seq data, whereas the blue peaks reflects the DRIP-sequenced data obtained from s9.6 immunoprecipitated HEK293T cells. The orange curve indicates Gro-seq data from HEK293T cells to quantify nascent RNAs at L1. The depicted schematic shows the L1 regions: 5’UTR, ORF1, and ORF2 (represented by black bars). The data also provides insights on the genomic locations of L1 in chromosome 7 of the hg38 assembly.

### p53 prevents the formation of L1 trans R-loop derived RNA-DNA hybrids

During the retrotransposition life cycle, the full-length L1 undergoes a process in which it generates an identical DNA copy of itself, known as L1cDNA, through the reverse transcription of L1mRNA. Hence, within the retrotransposition life cycle, there is potential for the formation of two types of RNA-DNA hybrids that can serve as R-loop; cis R-loops (L1mRNA-gDNA) occurring at the L1 5’-UTR promoter level during transcription, and trans R-loops (L1mRNA-cDNA) formed during reverse transcription. Genome-wide DRIP-seq studies reveal that cis R-loops are predominantly found in GC-rich regions and promoters ^42,43^. The recent findings demonstrate that p53 plays a role in global regulation of cis R-loops and viral proteins induce R-loops in p53 inhibited cells ^44,45^. However, the precise mechanism by which p53 regulates L1 derived R-loops has yet to be explored. The analyses of DRIPseq data from IMR-90, HEK293T, and k562 cell lines known to harbour p53 mutations (PMID: 26579211) revealed that young L1 subfamilies, which are capable of retrotransposition, participate in R-loop formation (**Figure** 2A). Further, to investigate whether L1 hyperactivation leads to increased accumulation of R loop formation, we treated A375 wt and p53^-/-^ cells with an HDAC inhibitor Sodium Butyrate (NaB) **(Figure 2B)**. RNAseq analyses on NaB (PMID: 38410440, 38410440) and TSA (PMID: 37496675) exposed cells and qRT-PCR on NaB treated A375 wt and p53^-/-^ did, in fact, show that L1 expression was elevated upon HDAC inhibition (**Figures** 2B and C). IF assays on wt and p53^-/-^ cells treated with NaB revealed an increased signal of s9.6 detecting the RNA-DNA hybrids (**Figures** 2D-2E’). Significantly, rescue experiments employing 3TC, a well-known reverse transcriptase inhibitor that inhibits L1cDNA formation, led to decreased s9.6 signals in both NaB-treated wild-type (wt) and p53^-/-^ cells. This implicates that L1 reverse transcription is involved in generating L1 R-loops (L1mRNA-cDNA) derived RNA-DNA hybrids (**Figures** 2E and E’). Subsequent qRT-PCR analysis of RNA-DNA hybrids precipitated by s9.6 antibodies from A375 wt and p53^-/-^cells, upon 3TC treatment, demonstrated complete depletion of R-loops from the L1-ORF2 region, with a modest reduction observed at the 5’-UTR (**Figure** 2F). This verifies the likelihood of L1-ORF2 being prone to trans R-loop formation. Similar outcomes were obtained via immunoprecipitation using a GFP antibody from cells expressing RNH-GFP (**Figure** 2G). These findings underscore that although the majority of the L1 5’-UTR region contributes to cis R-loop formation, L1-ORF2 likely facilitates trans R-loop formation, highlighting the involvement of L1 in both types of R-loop signals, arising from L1mRNA-gDNA as well as L1mRNA-cDNA hybrids.

**Figure 2.**
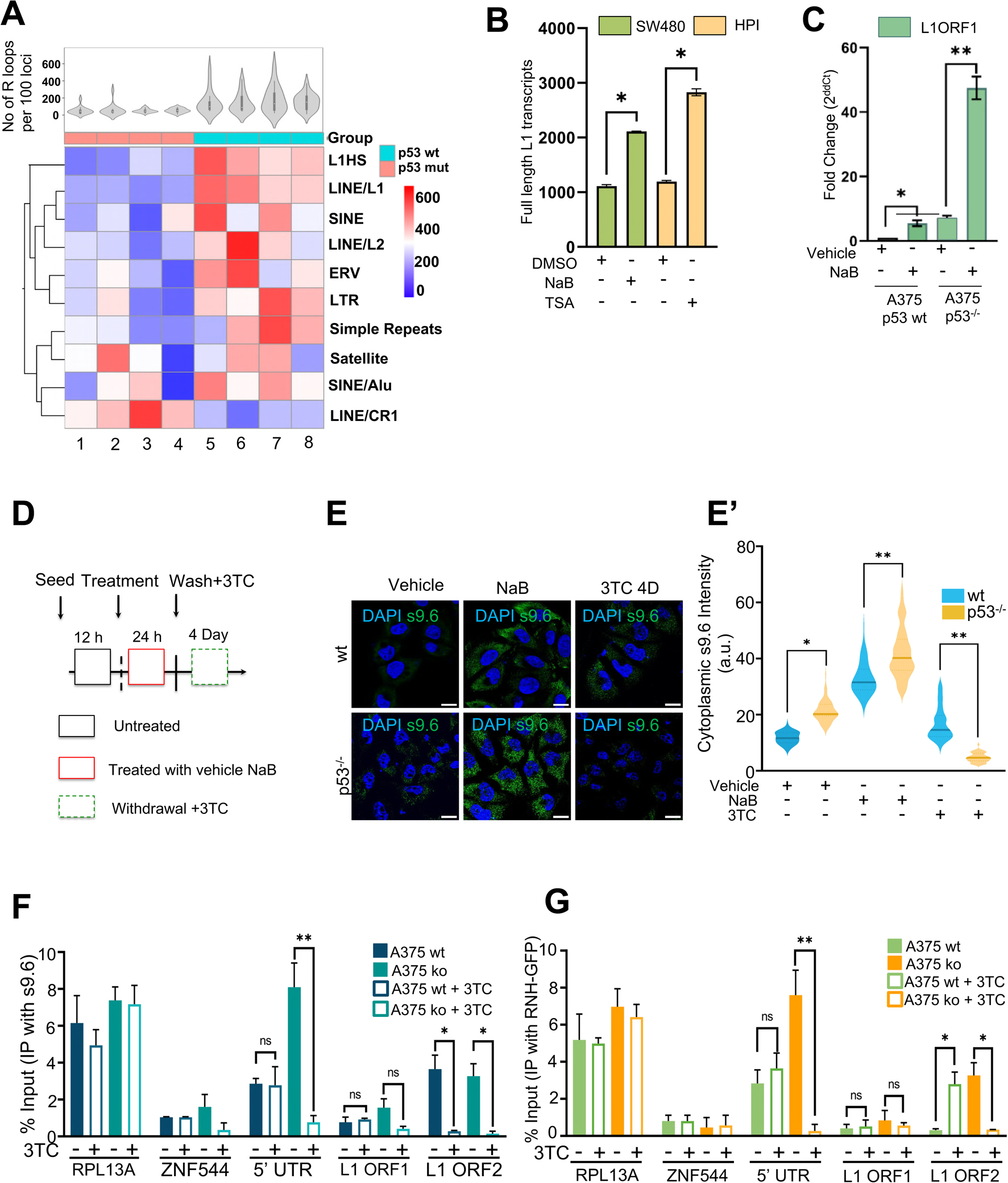
Increased accumulation of L1 trans R-loops derived RNA-DNA hybrids in p53^-/-^. (A) Heat map shows R-loops enrichment in human TE families obtained from the DRIP-seq data analysis. Each row represents a TE family grouped according to their repBase classification. The L1 has been classified as per their age, e.g. L1-young are full length sequences and are capable of retrotransposition including the loci from L1Hs family. Rows are clustered using the K-means algorithm and columns represent the sample types (with p53 status of the cells). The legend represents the differential presence of various TE families in the sample. Adjusted p-value of the Table is 0.05. Sample 1 (SRR2028291), 2 (SRR2028292), 3 (SRR5137197), 4 (SRR5137200), 5 (SRR2028293), 6 (SRR2028294), 7 (SRR9937575), 8 (SRR9937576). (B) Full-length L1 transcript analyses were performed in SW480 cell line (green) treated with NaB and from human pancreatic islets (HPI, yellow) treated with TSA (PMID: 38410440, 38410440). (Mann-Whitney U Test, *p<0.05). These cell lines are reported to harbour p53 mutation. (C) qRT-PCR analysis of L1 ORF1 expression of A375 wt and p53^-/-^ upon NaB treatment. The data shown are the averages of three biological replicates (Mann-Whitney U Test, *p<0.05, **p<0.01). (D) Illustration of strategies involving DMSO or NaB drug (25 µm) treatment on A375 wt and p53^-/-^, as indicated for 24 hours, followed by drug withdrawal for 4 and 7 days. Cell viability was assessed through cell glow titre experiments at (1 day, 3 days and 7 of drug treatment) (**Supplemental Figure S3A**) to measure HDACi drug toxicity. (E) Representative immunofluorescence microscopy images of three biological replicates showing RNA-DNA hybrid accumulation in wt and p53^-/-^ A375 cells after treatment with either DMSO or NaB for 24 hours, followed by 3TC treatment for 72 hours. Scale bar indicates 50 um. (E’) Quantification of s9.6 intensities from the whole cells was done using Fiji software ^62^. Violin plots exhibit combined data from three biological replicates with p values determined by Mann–Whitney test. p<0.05,*p<0.01. A minimum of 50 cells were measured in each sample of the three biological replicates. (F) DRIP-qPCR was performed to measure R-loops in A375 p53 wt and p53^-/-^ cells upon 3TC treatment. Immunoprecipitation carried out using the s9.6 antibody. Enrichment of RNA-DNA hybrids in each region, normalized to input values. Data represent mean±S.D. from three independent experiments. *p<0.05, ns=non-significant (Mann-Whitney U test). (G) DRIP-qPCR on RNaseH D210N-GFP stably transfected cells of A375 p53 wt and p53^-/-^ cells upon 3TC treatment. Immunoprecipitation carried out using the GFP antibody. Enrichment of RNA-DNA hybrids in each region, normalized to input values. Data represent mean±S.D. from three independent experiments. *p<0.05, ns = non-significant (Mann-Whitney U test).

Next, we conducted overexpression of L1-encoded proteins-ORF1p, EN and RT enzymatic domains of ORF2p c-terminally fused with GFP and empty vector (EV) as a negative control, in p53^-/-^ cells of A375 and U2OS cell to investigate the potential impact of these overexpressed proteins on the abundance of RNA-DNA hybrids. It has been established that the RT domain maintains reverse transcriptase activity autonomously within the cytoplasm ^24^. Overexpression of RT domain resulted in elevated levels of cytoplasmic RNA-DNA hybrids, consistent with recent findings reported by Kathleen’s group ^24^, as evidenced by increased s9.6 signals in the IF experiments (**Figure** 3A). This was accompanied by elevated transcript levels of DNA damage indicators (RAD51, BRCA1, and Rb) and inflammatory gene markers (IL6, STING pathway, IRF-7, MCP-1, CXCL-10, IFN-β, and ISG-15) measured using the qRT-PCR experiments (**Figure** 3B). Upon 3TC treatment, the augmented RNA-DNA hybrids in p53^-/-^ overexpressing ORF1p-GFP, EN-GFP, and RT-GFP were mitigated (**Figure** 3A’), confirming the involvement of L1cDNA derived RNA-DNA hybrid formation. This was corroborated by decrease in accumulation of RAD51, BRCA1, RB and inflammatory gene markers (**Figure** 3B) suggesting the aberrant formation of L1mRNA-cDNA with deleterious effects on genome stability. Moreover, to confirm if the overexpression of the RT-GFP leads to the generation of L1 trans R-loops, we conducted sequential DRIP-qPCR experiments (**Figure** 3C). These experiments involved immunoprecipitation using GFP from *in vivo* RNH-GFP expressing p53^-/-^ followed by IP with s9.6 antibodies to capture R-loops. This sequential immunoprecipitation assays validated that RT domain alone can specifically facilitate formation of R-loops in p53^-/-^. Following this, IF assays utilizing the yH2AX antibody demonstrated increased DNA damage in cells with overexpression of the RT domain (**Figures** 3D and D’) suggesting that RT domain alone is capable of inducing L1cDNA potentially culminating in the formation of L1trans R-loops derived RNA-DNA hybrids and subsequent DNA damage.

**Figure 3.**
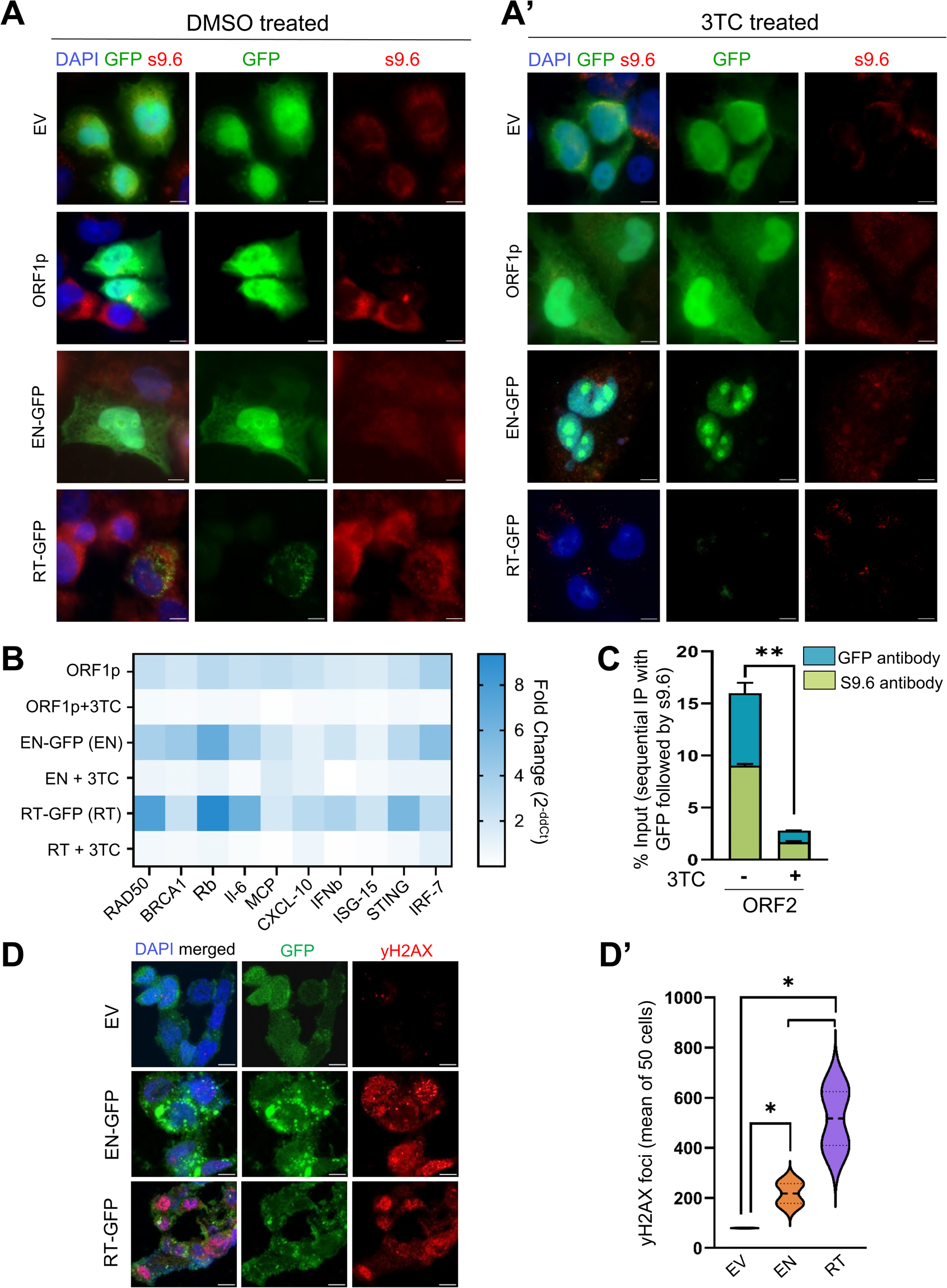
Effect of reverse transcriptase inhibitor on RT domain overexpressing p53^-/-^ cells. (A) Representative fluorescence images depict the transient overexpression of A375 p53^-/-^ cells with pEGFP-N1 (EV), L1-ORF1p, Endonuclease (EN), Reverse Transcriptase (RT) domains of L1-ORF2p followed by 3TC treatment for 3 days post transfection of 24 h. The overexpressed proteins are c-terminally GFP tagged (green) and s9.6 (red) that visualize RNA-DNA hybrids are mostly localised at the cytoplasm, nucleus is stained with DAPI (blue). Scale bar represents 50um. (B) The heatmap shows a consolidated qRT-PCR of DNA repair genes (RAD50, BRCA1, Rb) and inflammatory markers (STING, IRF7, MCP, CXCL10, IFNβ, ISG15) from the exogenously expressed ORF1p, EN domain, RT domain in A375 p53^-/-^ cells followed by 3TC treatment for 3 days. The subsequent legend shows the level of expression following overexpression. The samples were harvested after 3TC treatment for 3 days on 24 h transfected cells. The data from three biological replicates were included for heatmap generation. The normalised p-value of the data is less than 0.05. (C) Sequential DRIP-qPCR was performed on RT-GFP exogenously overexpressing A375 p53^-/-^ followed by DMSO or 3TC treatment for 3 days. The first round of immunoprecipitation was performed using GFP antibody followed by another round of immunoprecipitation by s9.6 on the GFP immunoprecipitated samples. Enrichment of RNA-DNA hybrids in each sample, normalized to the respective input values. Data represent mean±S.D. from the three independent experiments. (D) Representative confocal images of EV, EN, RT (all GFP tagged) transiently overexpressing p53^-/-^ cells (green) and yH2AX foci (red) localised at the nucleus (DAPI, blue). The merged images show overlap of yH2AX, DAPI, GFP. Scale bar represents 50um. (D’) Violin plots represent the y2AX (red) foci quantification. yH2AX foci were quantified via Fiji software. The mean (black line) represent the average foci from 50 nuclei. The data was quantified using fiji between three biological replicates. *p<0.05 (Mann-Whitney U test).

### p53 promotes establishment of L1 re-silencing

Given our findings that delineate trans-repressive functions of p53 in maintaining repression of L1 RNA-DNA hybrids, we sought to investigate its potential role in mediating the re-establishment of L1 silencing. To assess this, we employed HDACi drugs; TSA and NaB to induce L1 activation in a reporter system where 5’UTR-eGFP of L1 retrotransposons is inserted at the huAAVS1 site in both p53 wt and p53^-/-^cells of U2OS (**Figures** 4A-F) and A375 **(Figures** S2A-G) ^5^. Following a 24-hour treatment with HDACi drug (Figure 4A), activation of L1 reporters were assessed by quantifying the GFP positive cells by comparing the DMSO and 24 h TSA panels using the confocal microscopy. The efficacy of both wt and p53^-/-^ cells to suppress L1-5’UTR-eGFP upon drug withdrawal of 3 and 6 days elucidated a complete silencing of GFP in wt but not in p53^-/-^cells, this can be observed by comparing the panels of cells treated with DMSO and those observed at 4 and 7 days (**Figure** 4B). Similarly, the expression levels of egfp transcripts correlated with increased GFP intensities in L1-5’UTR reporter cells in response to HDACi **(Figures** 4C and 4D). This activation of GFP closely corresponded to activation of endogenous L1-ORF1, as demonstrated through RNA *in situ* hybridization using stellaris probes **(Supplementary Table 2 for probe sequences) (****Figure** 4B). Four days after the withdrawal of TSA, the activity of the L1-5’UTR reporter returned to uninduced levels in p53 wt cells, but not in p53^-/-^. No variation was detected in cellular proliferation using the cell glow titre test between wt and p53^-/-^ A375 L1-5’UTR-eGFP reporters following HDACi treatment **(Figure** S2A). Similarly, using qRT-PCR and microscopy, the reactivation of GFP and L1-ORF1 and subsequent re-silencing were also detected in A375 wt after 24 treatment with HDAC inhibitors followed by 4 days of withdrawal, but not in the p53^-/-^cells **(Figures** S2B-E). In line with previous research on HDACi treated embryonic stem cells ^16^, our experimental model demonstrated that inhibiting HDAC with TSA led to hyperacetylation of histones H3 and H4. This was evident from the increased signal observed in western blotting when probed with antibodies targeting H3Ac and H4Ac in both U2OS L1-5’UTR **(Figures** 4E and 4F) and A375 L1-5’UTR reporter cell lines **(Figures** S2F and S2G).

**Figure 4.**
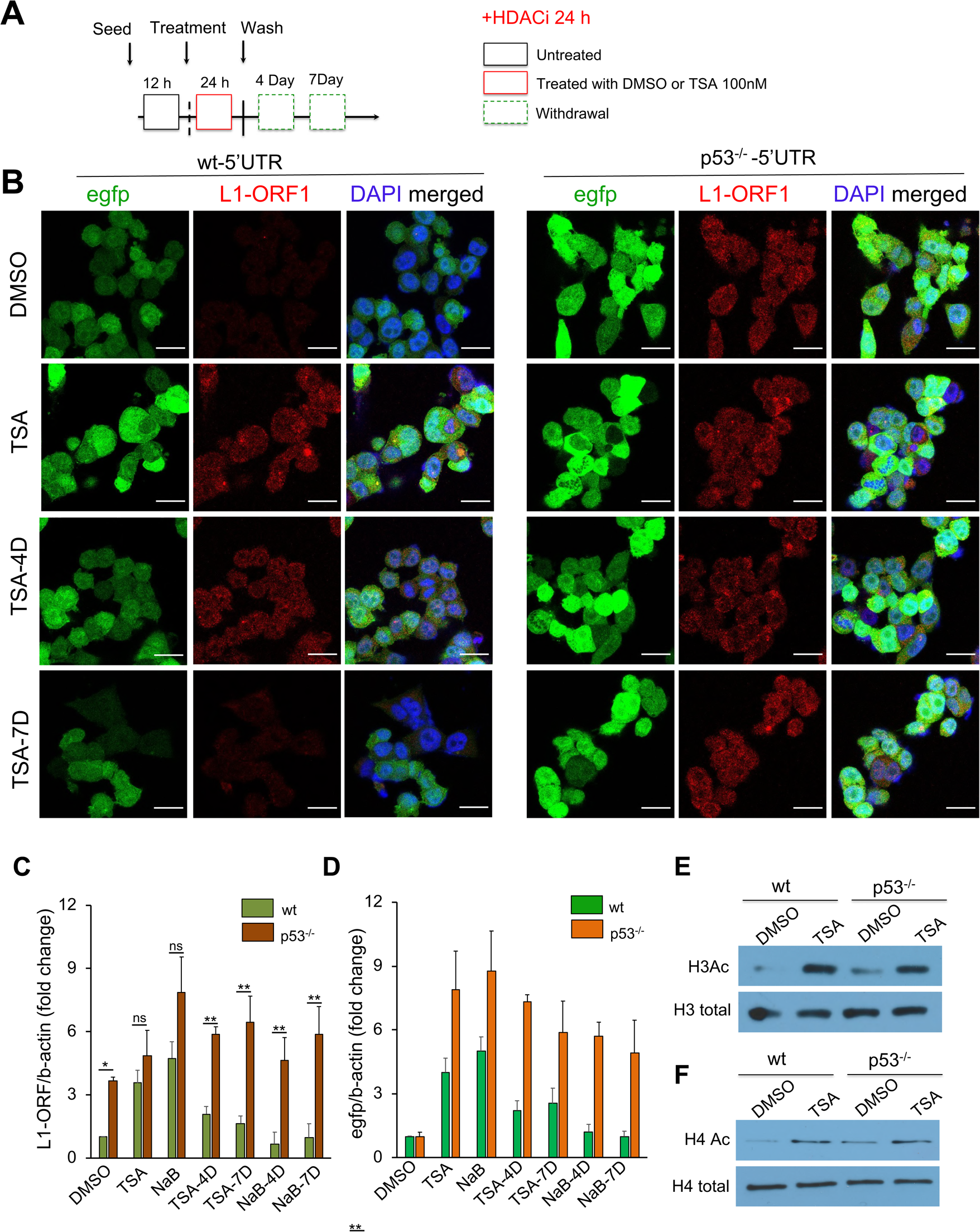
The dynamics of the L1-5’UTR-eGFP reporter upon exposure to HDAC inhibitors and subsequent drug withdrawal. (A) Illustrates the schematic of TSA drug treatment and withdrawal in U2OS wt and p53^-/-^ cells. The samples were collected for qRT-PCR, western blotting, and microscopy at 24 hours post-drug (HDACi) treatment, as well as at 4 and 7 days. TSA and NaB denote drug treatment for 24 hours, TSA-4D and NaB-4D indicate drug withdrawal of 3 days, and TSA-7D and NaB-7D signify withdrawal after 6 days post 24-hour treatment (L1-ORF1; 568, GFP; 488, and DAPI channels). (B) Confocal imaging of GFP and RNA *in situ* hybridization of L1-ORF1 using Stellaris probes (from LGC Biosearch Technologies) was performed in wt and p53^-/-^ U2OS cells harbouring L1-5’UTR-eGFP construct. The reporter cells were treated with either DMSO or TSA for 24 hours, followed by withdrawal for 3 and 6 days. Samples were collected at 1 day, 4 days, and 7 days for confocal imaging post HDACi treatment for GFP activation and silencing. Presented here are representative confocal images from three independent immunofluorescence experiments. Activation of L1-ORF1 transcripts was visualised using Qusar 570 Stellaris probes sequences (**Supplementary Table 2**). Scale bar indicates 20 um. (C-D) RT-qPCR analysis was performed to quantify L1-ORF1 and gfp transcripts in wt and p53^-/-^ L1-5’UTR cells treated with DMSO or TSA/NaB at 1, 4 and 7 days. The data represent the mean ± standard deviation (S.D.) from three independent biological replicates (n=3). *p<0.05, **p<0.01, ns=non-significant. (Mann-Whitney U test). The primers used for qPCR analysis of ORF1 and egfp transcripts are provided in **Supplemental Table 1.** (E-F) Western blotting of H3Ac and total H3 levels (E) and H4Ac and H4 total levels (F) in U2OS wt and p53^-/-^ cells upon either DMSO or TSA 24 h treatment.

**Figure 5.**
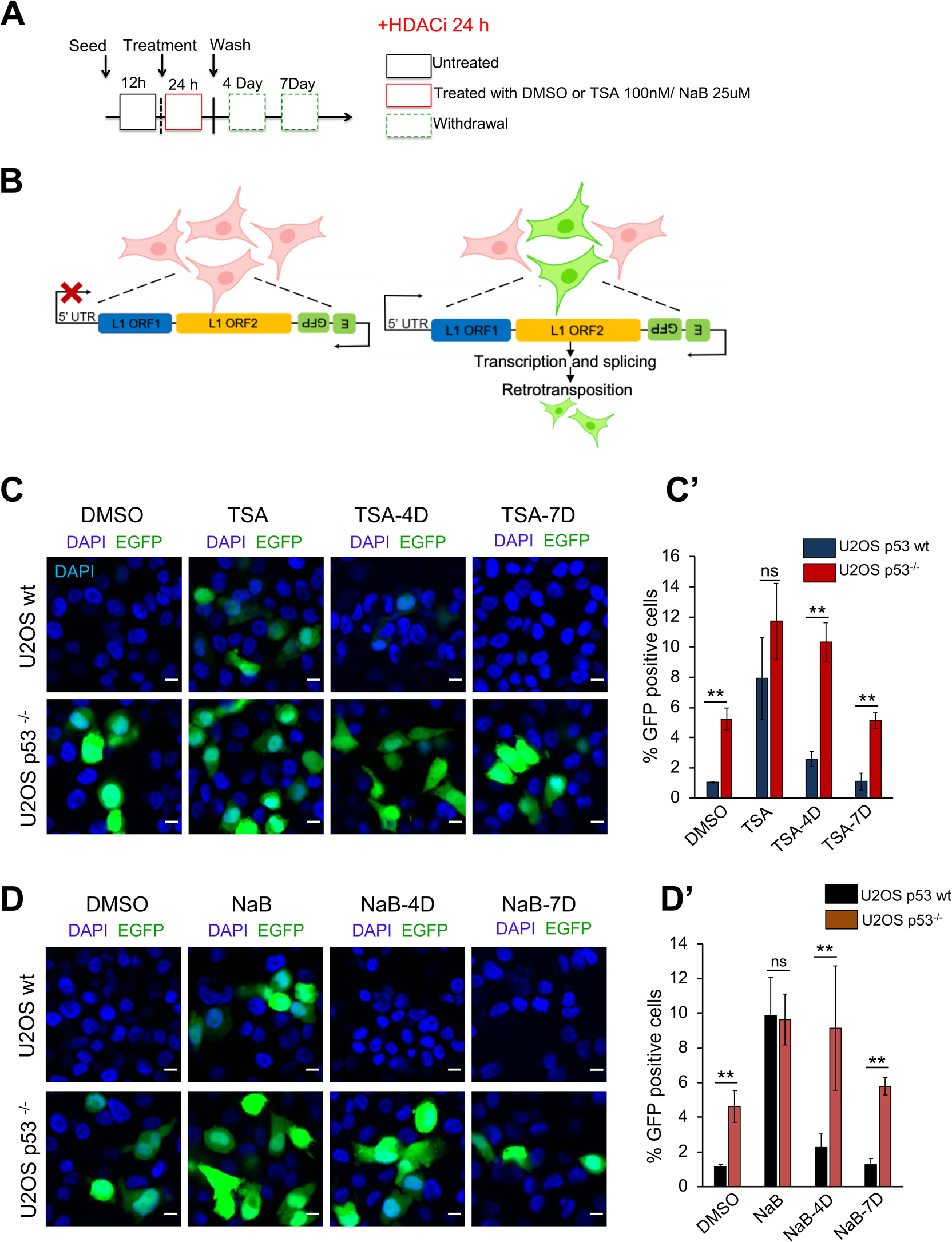
p53 facilitates the rapid re-establishment of LRE3 silencing, a retrotransposition indicator, following withdrawal of HDACi drugs. (A) Illustrates the schematic representation of DMSO, TSA/NaB drug treatment and subsequent withdrawal in U2OS wt and p53^-/-^ cells. (B) Illustrations indicates the nature of LRE3 plasmid which is a surrogate marker of retrotransposition. Cells that undergo retrotransposition are GFP positive (green). (C-D) The representative confocal images demonstrate the expression of GFP (green) in U2OS wt and p53^-/-^ cells following exposure to TSA (C) and NaB (D) drugs for 24 hours, followed by drug withdrawal of 3 or 6 days. Scale bar indicates 20um and the experiments were conducted with 3 biological replicates. (C’-D’) Percentage of GFP positive cells in U2OS wt and p53^-/-^ at 24 hrs of TSA (C) and NaB (D) treatment followed by drug withdrawal at 3 and 6 days. GFP positive cells were counted from minimum 10 fields from the 3 independent experiments. The error bar represents mean of 3 biological replicates. *p<0.05, **p<0.01, ns=non-significant (Mann-Whitney U test).

Next to determine whether p53 also enables resilencing of retrotransposition activation, we used a synthetic L1 designated 99-gfp-LRE3 retrotransposition indicator ^46,47^, using HDACi treatment, similar experiment was conducted on stably transfected parental wt and p53^-/-^ U2OS cells, schematic shown (**Figure** 6A). This indicator **Figure** 6B mimics retrotransposition cycle generating all L1 intermediate components allowing direct measurements of *de novo* retrotransposition rates by visualising GFP positive cells (green cells). Consistent with the L1-5’UTR reporter, we noted re-silencing of L1-retrotransposition in wt cells upon withdrawal experiments after TSA and NaB treatments. Whereas no re-silencing was observed in p53^-/-^ cells as quantified in (**Figures** 6C-6D’).

**Figure 6.**
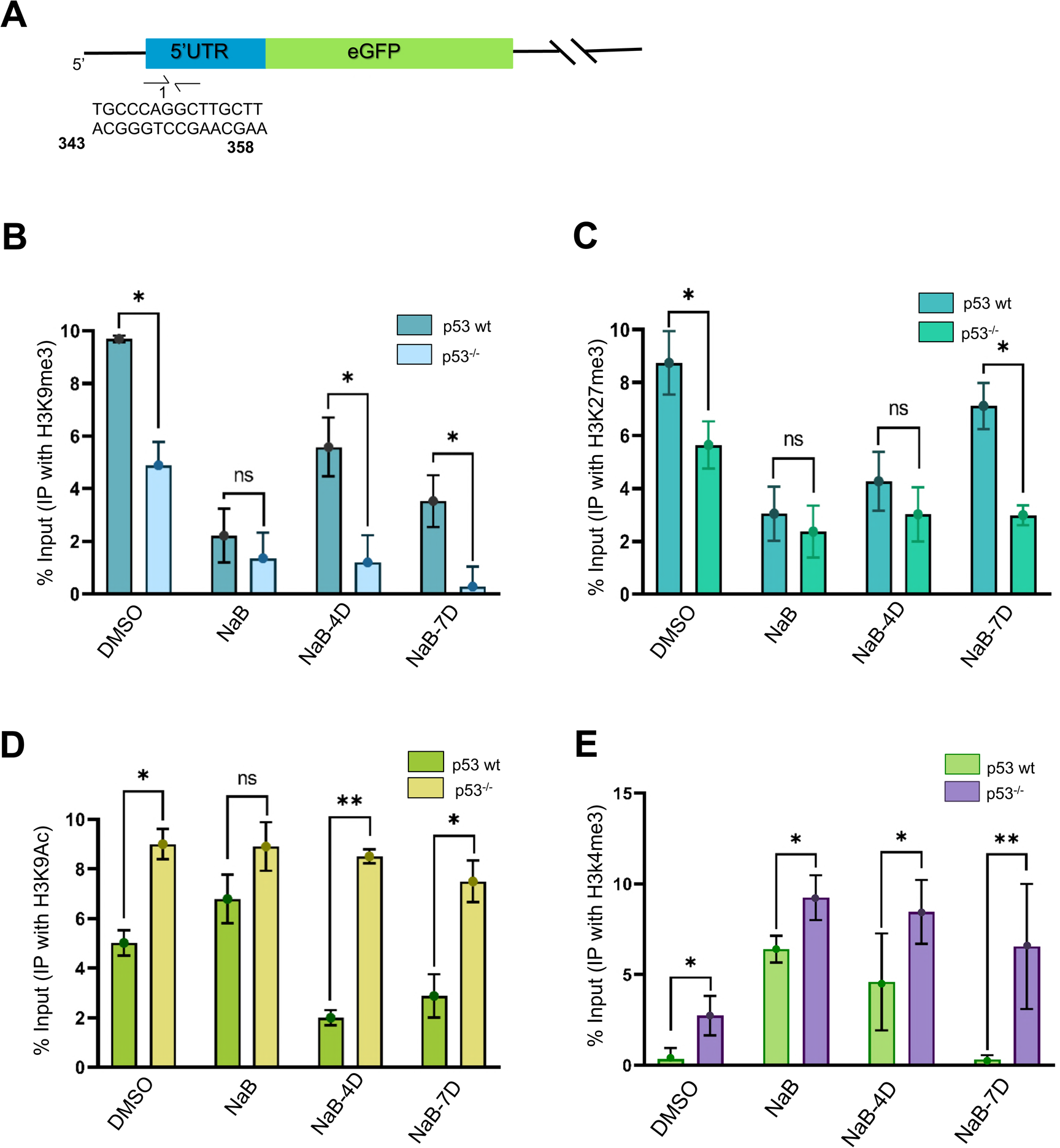
Histone modification dynamics in L1-5’UTR-eGFP reporter cells upon HDACi exposure and withdrawal. (A-C) Chromatin immunoprecipitation followed by qPCR analyses was conducted on the 5’UTR region of A375 wt and p53^-/-^ L1-5’UTR-eGFP stable cells. Primers targeting the p53 binding region were utilized, with immunoprecipitation carried out using antibodies against histone modifications H3K9me3, H3K27me3, H3K9Ac, and H3K4me3. The results illustrate normalized levels of H3K9me3 and H3K27me3, representative of repressive histone marks (A and B), normalized to input values, as well as levels of activation marks H3K9Ac (C) and H3K4me3 (D). Reporter cells were treated with HDACi for 24 h followed by withdrawal for 3 and 6 days. Samples for analyses were collected at 1 day, 4 and 7 days (for drug withdrawal), as depicted (refer to Figures 4 and 5). It is noteworthy that alterations in H3K9me3, H3K27me3, H3K9Ac, and H3K4me3 enrichment are specific to the p53 status. Error bars represent the standard error of the mean from three biological replicates. *p<0.05, **p<0.01, ns=non-significant (Mann-Whitney U test).

### p53 promotes restoration of deposition of histone repressive marks at L1-5’UTR

In a previous study, we demonstrated that p53 regulates L1 transposons by epigenetic mechanisms ^9^. To investigate whether p53 can facilitate re-silencing of L1 by recruiting these marks, we activated L1 transposons in both A375 wt and p53^-/-^ L1-5’UTR-eGFP reporters, as demonstrated in **Figure 6**. The quantification of histone repressive marks H3K9me3 and H3K27me3 was performed with ChIP-qPCR studies using primer sets targeting the p53 binding site at the L1-5’UTR (**Figure** 6A). Upon treatment of cells with HDACi drug (NaB), L1 activation coincided with depletion of H3K9me3 (**Figure** 6B) and H3K27me3 (**Figure** 6C), as well as acquisition of H3K9Ac (**Figures** 6D) and H3K4me3 (**Figures** 6E). Further, levels of H3K9 and H3K27 trimethylation reversed back to the DMSO treated levels following a 4-day drug withdrawal experiment in wt cells with a concurrent decrease in H3K9Ac marks and H3K4me3. Notably, changes were not observed in p53^-/-^ cells. Importantly, this alteration in chromatin marks closely correlated with the reversal of L1 induction observed in wt cells, providing the evidence that p53 initiates the re-silencing of activated L1 transposons by promoting the recruitment of epigenetic repressive marks and removal of activating marks (**Figures** 4, 6 and S2).

### A LINE1 reverse transcriptase antagonist partially reverses DNA damage triggered by L1

The findings depicted in **Figures** 4 and S2 demonstrate that p53 establishes resilencing of L1-transposons upon HDACi withdrawal by mediating recruitment of epigenetic repressive and inhibiting activating marks at L1-5’UTR. HDACi is shown to induce DNA double-strand breaks in cancer cells^48^. Therefore, to investigate the involvement of p53 in the reversal of L1 retroelement-induced DNA damage, we subjected U2OS wt and p53^-/-^ cells to HDACi (**as shown in** **Figure** 7A) (**as shown in Figure** S3), followed by exposure to either DMSO or 3TC. Previous study from us has shown L1-mediated genomic rearrangement in early passages of p53^-/-^cells ^4^, and similarly, we observed induction of L1-ORF1p de repression accompanied by DNA damage upon TSA treatment in both the wt and p53^-/-^ U2OS cells. The treatment of p53^-/-^ cells with 3TC resulted in restoration of L1-ORF1p expression to the vehicle (DMSO) treated level, accompanied by a notable decrease in the number of 53BP-1 foci, which serve as an indicator of DNA damage (**Figure** 7A). Repair of DNA damage and suppression of inflammatory genes to their uninduced levels occurred promptly in wt cells, while it was impaired in p53^-/-^ cells. Similar results were observed in staining with γH2AX, an additional marker of double-stranded DNA breaks. **(Supplemental Figure** S3 A and A’). Collectively, our findings suggest that p53 facilitates the suppression of activated L1 transposons that trigger DNA damage and inflammation. The immunofluorescence experiments on cells treated with DMSO and NaB to evaluate the impact of HDACi on the acetylation and methylation levels of H3K9 and H3K27 histones in both wt and p53^-/-^ cells detected a global increase in the acetylation of lysine residues on the H3 protein, notably at positions 9 and 27, accompanied by a reduction in methylation at the same lysine residues **(Figures** S4 and S5). Administering 3TC for 4 and 7 days led to the re-establishment of these histone marks in wt cells, but not in cells lacking the p53 gene.

**Figure 7.**
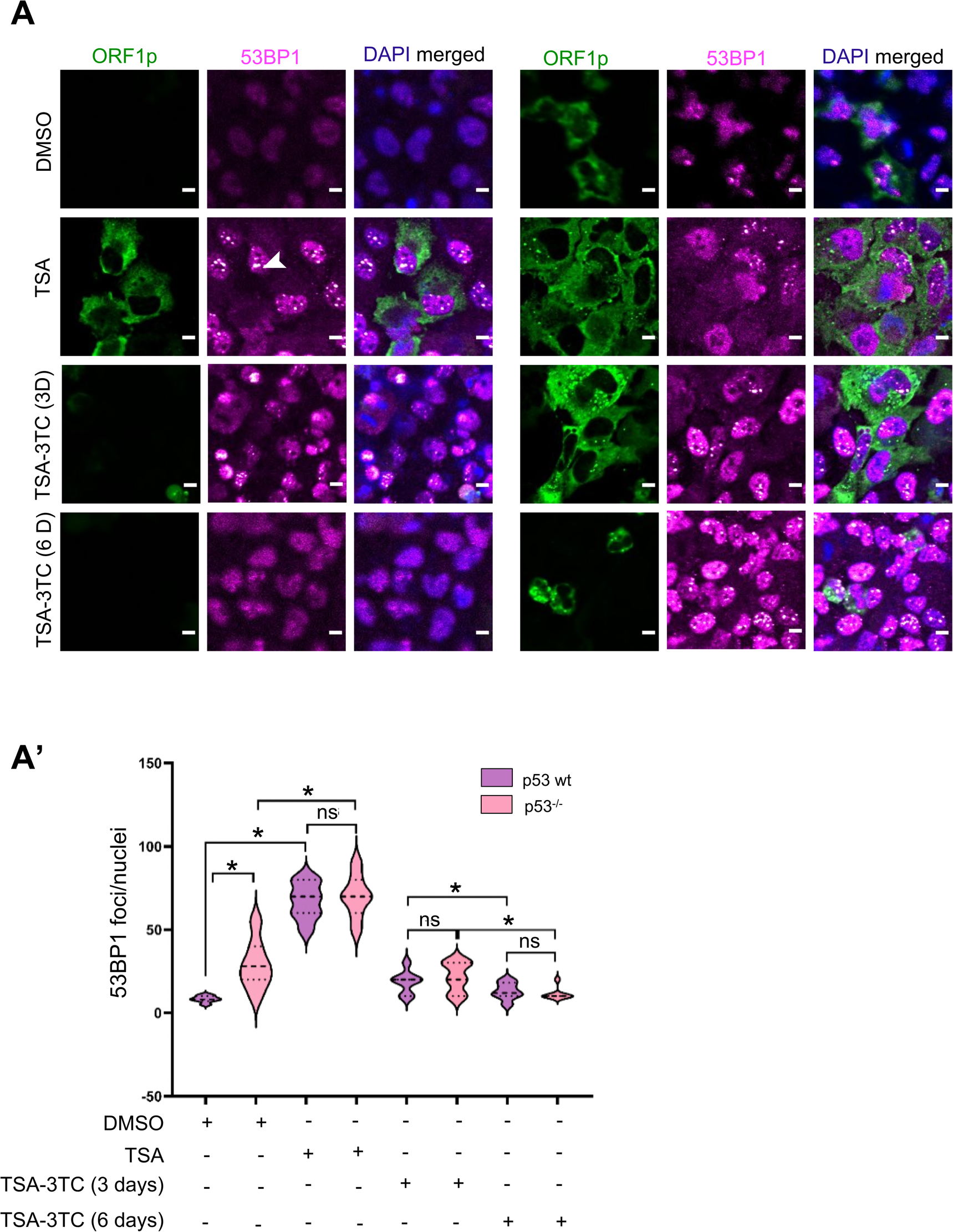
Effect of 3TC on L1-ORF1p expression and 53BP-1 foci. (A) Confocal imaging of L1-ORF1p (green), 53BP-1 (magenta, foci indicated in white) and nucleus (blue) in U2OS wt and lacking p53 were treated with either DMSO or TSA for 24 hours, followed by 3TC treatment for 3 and 6 days. Imaging of cells performed at 1 day after TSA exposure, and at 3 and 6 days after 3TC treatment. Presented here are representative confocal images from three independent immunofluorescence imaging experiments. Upon 24-hour exposure to TSA, activation of L1-ORF1p and accumulation of 53BP-1 foci (indicated by white arrows) were observed. Conversely, suppression of L1-ORF1p and reduction of 53BP-1 foci were more evident in wt than p53^-/-^ cells after 3 and 6 days of 3TC treatment. Scale bar denotes 20um. (A’) The Violin plot shows the quantified 53BP1 foci from microscopy images. The bold dotted line represent the mean foci from 50 different nuclei. The quantification has been done from 3 different biological replicates. *p<0.05, ns=non-significant (Mann-Whitney U test).

## Discussion

Our comprehensive analyses, combining experimental and bioinformatic approaches, unveil the pivotal function of p53 in repressing RNA-DNA hybrids originating from L1 retrotransposition derived R-loops encompassing both the cis (L1mRNA-gDNA) and trans (L1mRNA-cDNA) configurations **(see Figures** 1 and 2). While previous studies primarily focused on the mutagenic potential of L1 *de novo* insertions, our understanding of the impact of nucleic acid intermediates like L1 RNA-DNA hybrids and protein products such as L1-ORF1p and ORF2p in genomic instability and inflammation remains limited. Despite the relatively infrequent detection of new L1 insertions in cancer genomes compared to their deleterious activity, L1 activation is a common feature in many sporadic cancer genomes, as evidenced by recent pan-cancer studies ^25,49,50^. Our finding regarding the prominent involvement of L1 promoter in cis R-loop formation aligns with previous genome-wide studies highlighting the enrichment of R-loops in promoter regions globally ^41^. This is corroborated by the observation of overlapping peaks of DRIP-seq at the L1 promoter with those identified from the GRO-seq, suggesting the occurrence of transcriptional pausing at L1-5’UTR that leads to the formation of cis RNA-DNA hybrids. Conversely, the participation of L1-ORF2 genic region in trans R-loop formation is validated by our observation of diminished R-loop signals upon 3TC treatment in RNH-GFP expressing cells (**Figure** 3). R-loops are capable of yielding both adverse and beneficial physiological outcomes in a cell ^51^. This prompted our speculation that R-loops originating from retrotransposons could impede the finalization of insertions—a critical controlling step in the retrotransposition process to prevent the amplification of L1 copies in the genome. However, the observed decrease in DNA damage markers and inflammation upon RT overexpression, coupled with a reduction in cytosolic RNA-DNA hybrids upon treatment with 3TC in DRIP-qPCR and immunofluorescence assays, provides mechanistic insights into the contribution of RNA-DNA-derived L1 R-loops to eliciting genome instability and inflammation, the two hallmarks of cancer. This also indicates that, in addition to the canonical retrotransposition insertion, other intermediate components can have oncogenic potential. These observations led us to speculate that L1 R-loops may not only act as a bystander but as a double edged sword in this context.

Furthermore, the inhibition of HDACs resulted in the loss of methylation and the gain of acetylation at the 5’UTR, activating L1 transposons and the derived R-loops (refer to **Figures** 3, 4, 5, and S2). Withdrawal of HDACi caused permanent L1 de repression in p53^-/-^ cells, inducing DNA damage and inflammation triggered by L1 R-loop-derived RNA-DNA hybrids. In contrast, wild-type cells swiftly silenced L1 by promoting H3K9me3 and H3K27me3 (**Figures** 6 and 7). This mechanism may confer cancer resistance in p53-driven cancers treated with HDACi ^52^. Additionally, treatment with 3TC prevented the activation of L1 RNA-DNA hybrids representing the first evidence of trans L1 R-loop formation in p53^-/-^ unstressed cells as well as in HDACi-treated cells.

Our findings also suggest that p53 promotes re-establishment of silencing of hyperactive L1s, advancing our prior research underscoring the function of p53 in preserving L1 repression in human somatic cell lines. Considering the function of p53 in mediating repression of L1s and L1-derived R-loops, especially those originating from nuclear L1mRNA-cDNAs, these discoveries could hold significant implications for elucidating the mechanisms underlying p53 tumor suppression. Unlike conventional understandings, this study reveals that the RT domain is involved in regulating RNA-DNA hybrids that triggers DNA damage as seen in **Figure 3**. These unexplored oncogenic risks posed by the L1 RT domain trigger acute inflammatory transcriptional programs upon overexpression, particularly in the absence of p53 ^53–55^. Treatment with 3TC definitively prevented the induction of nearly all inflammatory genes and DNA damage (**Figure 3)** in both wt and p53^-/-^. Since 3TC is a potent inhibitor of the L1 reverse transcriptase ^56–58^, these results indicate that L1 retrotransposition intermediates provoke inflammatory programms in p53 deficient cells. Furthermore, these findings also suggest that the induction of immune response genes may stem from the RT domain of ORF2p, attributed to the cytosolic accumulation of RNA-DNA hybrids, along with the accumulation of trans L1 R-loops assessed in sequential DRIP-qPCR assays on RNH-GFP *in vivo* system. Taken together, these findings suggest that incorporating a reverse transcriptase inhibitor into personalized cancer therapy ^59^ could enhance the efficacy of HDAC inhibitor treatment for cancer.

## Materials and Methods

### Cell culturing

p53 wild-type (wt) and p53 knockout (p53^-/-^) cells of A375, U2OS cells were obtained from Prof. John Abrams, UT Southwestern Medical Center, Dallas, Texas. A375 and U2OS cells were cultured in Dulbecco’s Modified Eagle’s Medium (Gibco, #12100046). All cell lines were grown at 37°C, 5% CO2 and passaged using Trypsin (Gibco, #15400054) in the presence of an anti-antimycotic antibiotic (Gibco, #15240062) .

### Generation and screening of p53 knockout cells

CRISPR cas9 editing (pX458), was used to generate p53^-/-^ cell lines in the laboratory of Prof. John Abrams at UTSW. p53^-/-^ clones were identified by western blot using p53 monoclonal DO-1 (sc-126) and L1-ORF1p expression levels were examined using anti L1-ORF1p antibody using immunofluorescence experiments. Individual clones were grown until 100% confluent in 6-well plates and whole cell lysates were prepared in RIPA lysis buffer (NaCl 140mM, EDTA 1uM, Tris HCl 10mM, Triton X-100 1%, SDS 0.1%, Sodium Deoxycholate 0.1%) containing protease inhibitors (Roche). Protein was quantified using Bradford assay (Protein Assay Kit, BioRad) and an equal amount of total protein was loaded. Antibodies used were as follows: p53 monoclonal DO-1 (sc-126), anti L1-ORF1p, clone4H1 (Millipore, #MABC1152) and β-actin (sigma, #A-2228). Sanger sequencing was performed to confirm p53^-/-^.

### Generation of huAAVS1 targeted L1-5’UTR-eGFP reporter cells

The wt L1-5’UTR constructs, previously generated and kindly provided by Prof. John Abrams at UTSW, were utilized. Stably transfected cells were created using huAAVS1-TALENL (Addgene #59025) and huAAVS1-TALENR (Addgene #59026), along with donor plasmids. Three days after transfection, single cells positive for GFP were sorted into 96-well plates using fluorescence-activated cell sorting (FACS), following the protocol outlined in ^4^.

### For 3TC experiments

2.5 x 10^5^ wt and p53^-/-^ A375 and U2OS cells were plated into 6-well plates. After overnight of seeding media was changed with fresh media containing DMSO, HDACi. After 24 h of HDACi treatment media was replaced with fresh containing 3TC (10 µM). After 4 days and 7 days of 3TC treatment, cells were harvested for RNA isolation or analysed for microscopy. qRT-PCR was conducted using Applied Biosciences QuantStudio 5. Gene expression was normalized to β-actin, and fold induction was calculated relative to the wild type. Primer sequences utilized in this study are available in **Supplemental Table 1**.

### Real Time PCR experiments (RT-PCR)

Total RNA was isolated using TRIzol (Invitrogen), then treated with Turbo DNase (Thermo Fischer, #AM1907) for removal of DNA. RT-PCR was conducted utilizing cDNA synthesis kit Verso (Thermo Fischer, #AB1453A). Applied biosciences Quant studio 5 was used for real time PCR and normalization of gene expression was done against β-actin (for standard gene expression analysis), RPL134 for DRIP-qPCR experiments, and fold induction was determined relative to the wild type. Refer to Supplemental Table 1 for the primer sequences employed in RT-PCR assays.

### Effect of HDACi in wt and p53 cells expressing LRE3 retrotransposition

2.5 x10^5^ cells were transfected with equal amounts of 99-GFP-LRE3 (LRE3) plasmid, as previously described [34]. After 72 h, cells were selected for puromycin resistance. Similar protocol was followed for HDACi exposure followed by 3TC as mentioned in **Figure** 4. Confocal imaging was performed using LSM780 and LSM880 at the UTSW, Texas. GFP positive cells were counted from 5 images from each of the samples. 2 biological replicates were performed.

### Statistical Analysis

Quantitative data were collected from experiments conducted in at least two or three biological replicates with technical replicates. One tailed Student’s t-tests were calculated from three biological replicates to detect statistically significant differences (with a cutoff of *p<0.05 and **p<0.01).

### Drug treatment

TSA (Sigma Aldrich, #T8552) and NaB (Sigma Aldrich, #567430) were purchased Sigma. All drug treatments were carried out in DMEM (Gibco, #12100046) supplemented with 10% (vol/vol) FBS (Gibco, #A5256701) and 100X anti-antimycotic (Gibco, #15240062). Treatment with TSA (100nM) was for 24 hr as indicated in the figure legend. NaB was used at 25 μM as indicated.

### Cloning of NLS tagged RNaseH1 with D210N

For NLS-tagged RNaseH1D210N, cDNA fragments (purchased from IDT) ^60^ containing human RNaseH1 with D210N catalytic dead mutant was amplified by PCR for cloning in pEGFP-N1 using EcoRI (NEB, #R3101S) and KpnI (NEB, #R3142S) restriction sites. One SV40 nuclear localization signal (NLS) (CCCAAAAAGAAACGCAAAGTG) was introduced by forward primer at the PCR step. Primers used in PCR amplification are listed in **Supplemental Table 1**.

### Generation of stable cell lines expressing RNaseH1 D210N (RNH-GFP)

To study R-loop in wt and p53^-/-^cells, nucleus localization sequences containing GFP-tagged, catalytically-inactive RNase H1 D210N was cloned in pEGFN-1. The cloned sequence was verified by sanger sequencing and A375 wt and p53^-/-^ cells were transfected for the stable cell line generation using G418 (Gibco, #11811064).

### ChIP-qPCR

One billion A375 wt and p53^-/-^cells were grown in a 150 cm^2^ dish and were fixed with 1% formaldehyde in flasks at room temperature for 10 minutes and subsequently washed with ice-cold PBS before a 125mM glycine solution was added to stop the fixation. The cells were scraped from the flask and were resuspended in ChIP lysis buffer (50 mM HEPES-KOH pH7.5, 140 mM NaCl, one mM EDTA pH8, 1% Triton X-100, 0.1% Sodium Deoxycholate, 0.1% SDS, Protease inhibitor). The cell lysate was sonicated using an M220 Focused-ultrasonicator (Covaris) to release chromatin lengths to 400bp, peak power 75.0, Duty factor =10, cycles/Burst =200, for 240 seconds. Chromatin was diluted ten times with RIPA buffer and incubated with the following antibodies – H3K9me3 (Cell Signalling Technologies #13969), H3K27me3 (Cell Signalling Technologies, #9733), H3K9Ac (Cell Signalling Technologies, #9649) . Parallel protein A/G beads (EMD Millipore, #IP0515ML), H3K4Me (Cell Signalling Technologies, #9751) were charged and incubated with the antibodies overnight, following high salt, low salt, and LiCl washes the next day and eluted. Eluted chromatin was treated with RNAase A (Thermo Fischer, #12091021) and proteinase K (NEB, #P8107S), crosslinks were reversed, and samples were purified using a PCR purification kit (Qiagen). ChIP enrichment was quantified by qPCR with Quant Studio 5 (Applied Biosciences).

### DRIP-qPCR

One billion A375 p53 wt and p53^-/-^ cells or cells expressing RNH mutant were plated into a 150 cm^2^ dish for overnight (Tarsons) and were fixed by 1% formaldehyde, followed by quenching via Glycine to a final concentration of 125 mM. Cells were rinsed twice with 10 mL cold PBS. 5 mL of cold PBS was added, dishes were scraped thoroughly with a cell scraper and transferred. Cells were lysed in 750 ul lysis buffer composed of 140 mM NaCl, 1 mM EDTA (pH8),1% Triton X-100, 0.1% Sodium Deoxycholate,0.1% SDS, Protease Inhibitors (added fresh each time) per 10 million cells. Cell lysis was performed at room temperature. The cell lysate was sonicated to shear DNA to an average fragment size of 400 bp by the M220 Ultrasonicator (Covaris) as previously mentioned for ChIP. Approximately 25 μg of RNA-DNA hybrid per IP was used. Each sample was diluted 1:10 with RIPA Buffer (50 mM Tris-HCl, pH8,150 mM NaCl, two mM EDTA pH8, 1% NP-40, 0.5% Sodium Deoxycholate, 0.1% SDS, Protease Inhibitors (added fresh each time). Primary antibody RNA-DNA hybrid (s9.6) antibody (Merck Millipore, #MABE1095, 3ul) or GFP (Thermo Fisher, 5ul) was added to all samples and incubated at 4°C for 2 h. 60 ul of Pierce A/G Agarose beads were taken for 25 ug of RNA-DNA hybrid. Beads were charged and incubated with the antibody incubated sample at 4 degrees overnight in shaking. The immunoprecipitated samples were washed by low salt, high salt, and LiCl wash buffers. RNA-DNA hybrids were eluted by slowly adding 120 μL of elution buffer (1% SDS, 100mM NaHCO3) to the protein A/G beads and vortex for 15 min at 30°C. 4.8 µL of 5 M NaCl and 2 µL RNase A (10 mg/mL) were added and incubated while shaking at 65°C overnight. 2 µL proteinase K (20 mg/mL) was added and incubated while shaking at 60°C for one hour. The RNA-DNA hybrids were purified using a PCR purification kit (QIAGEN). Analysis of RNA-DNA hybrid level was performed by qPCR.

### Quantification of RNA-DNA hybrids

RNA-DNA Hybrid levels were measured by RT-PCR using 2X POWER UP SYBR Green (Invitrogen, #A25741) and analyzed via the QuantStudio 5 Real-Time PCR System (Applied Biosciences). Primer sequences are listed in **Supplemental Table S1**. The qPCR results were analysed using the percentage input method. The RNA-DNA hybrid enrichment was calculated based on the IP/Input ratio. Three biological replicates were performed. The result is the average of all the biological replicates.

### Immunofluorescence

A375 p53 wt and p53^-/-^ cells were cultured on poly-L-lysine (Sigma, #P4707)-coated coverslips, washed with PBS and fixed with 4% paraformaldehyde. The cells were treated with 0.1% Triton X-100 in PBS and 10% fetal bovine serum and were subsequently incubated with primary antibodies at room temperature for 1 hour or 4 degrees overnight. After washing, secondary antibodies were added and incubated in the dark at room temperature for 1 hour. Cells were washed with PBS and counterstained with DAPI before observation under an Apotome microscope (Zeiss). All images were taken using the same microscope parameters, and fluorescence quantification was performed using fiji J software. Primary antibodies used were LINE1 ORF1p (Merck Milipore, #MABC1152), Anti RNA-DNA Hybrid antibody (Merck Milipore, #MABE1095), yH2AX (Invitrogen, #MA1-2022), H3K9me3 (Cell Signaling Technologies, #13969), H3K27me3 (Cell Signaling Technologies, #9733), H3K9Ac (Cell Signaling Technologies, #9649), and 53BP1 (Cell Signaling Technologies, #4937), all at a dilution of 1:50. Secondary antibodies used were: anti-mouse 568 (Thermo Fischer, #A-11031), anti-mouse 488 (Thermo Fischer,#A-10680), anti-rabbit 488 (Thermo Fischer, #A-11008), and anti-rabbit 568 (Thermo Fisher, #A-11011), at a dilution of 1:1000.

### Microscopy

Images were captured using 60x oil objective Zeiss LSM780, LSM880 (UT Southwestern Medical Center, Dallas, Texas) and Leica sp8 (ILS Bhubaneswar BT/INF/22/SP28293/2018) confocal microscope. Zeiss Apotome 3 fluorescent microscope (IISER Berhampur) was also instrumental for imaging. All images were processed via Fiji software package.

### Cytotoxicity assay using cell glow titre assay

A total of 5000 A375 wt-5’UTR and p53^-/-^ -5’UTR cells were distributed evenly into 96 well plates, with each well containing either DMSO or a solution of NaB or TSA followed by 3TC. The CellTiter-GLO kit from Promega (G7570) was utilized. Cells were treated with HDACi for 24, 48 or 72 h. 25 ul of CellTiter-Glo® Reagent was added and mixed thoroughly for 2 minutes using an orbital shaker. The plate was kept at room temperature for 10 minutes and the luminous signals were quantified using a plate reader after 24, 48 and 72 h of treatment as per the instructions. Results are quantified in relation to the DMSO control for each genotype.

### RNA *in situ* hybridization assays

LGC Biosearch Technologies’ Stellaris^®^ RNA FISH Probe Designer was used for the designing of Qusar-568 probes for L1-ORF1 probes, for RNA *in situ* hybridization. Protocol was followed as per the manufacturer’s instructions.

### Dataset Analyses

DRIP-seq data of various cell lines (PMID 26579211), were retrieved from the NCBI Gene Expression Omnibus (GEO) in SRA format. Before analysis, the data underwent preprocessing to remove adapters and low-quality bases. Subsequently, the cleaned reads were aligned to the GRCh38 human reference genome assembly (hg38) using the STAR aligner (version 2.7.10a). To identify regions enriched with potential R-loops, MACS3 (Model-based Analysis of ChIP-Seq) (version 3.0.0) with the ’macs3 callpeak -t’ option was employed, ensuring precise localization within the genome. Additionally, RepeatMasker (version 4.1.6) was applied, utilizing the ’RepeatMasker - engine rmblast’ option, to mask retrotransposable elements and repetitive sequences, thereby maintaining the integrity of subsequent analyses ^61^. Expression levels of retrotransposable elements were quantified from the alignment files generated by STAR, followed by normalization for library size and transcript length. Heatmaps illustrating the expression patterns of retrotransposable elements across different cell lines were generated using the R programming language, with visualization facilitated by the gplots package. Visualization of peaks across the full length of LINE1 was achieved through the UCSC Genome Browser.74.

## Acknowledgements

This work was supported by the DBT/Wellcome Trust India Alliance fellowship (IA/I/22/2/506501) to Bhavana Tiwari. Pratyashaa Paul was supported by IISER Berhampur, (A.K) CSIR-JRF, (A.K.D) UGC-JRF. The authors would also like to acknowledge the assistance of ILS Bhubaneswar confocal imaging facility and the imaging facility UTSW, Dallas, Texas for microscopy. We acknowledge Prof. John Abrams at the Department of Cell Biology, University of Texas Southwestern Medical Center, Texas for generously providing chemicals, reagents, including wt and p53 ko cells, as well as plasmid resources utilized in this study. We thank Dr. Amanda Jones and Prof. John Abrams of UTSW, Texas for valuable suggestions. Illustration work for graphical abstract was created with BioRender.com. We thank IISER Berhampur for the support.

## Contributions

BT conceived the studies. PP, AK, AKD, AP, GP, FP and BT performed experiments and analysed data. SK processed the microscopy data and assisted in the illustration of the graphical abstract. BT designed the study, analysed data, and both BT and PP wrote the manuscript.

