## Supplementary Information for "p53 mediated regulation of LINE1 retrotransposition derived R-loops"

Bhavana Tiwari, PhD
Intermediate Fellow, DBT-Wellcome Trust India Alliance
Department of Biological Sciences,
Indian Institute of Science Education and Research Berhampur,
Berhampur, Odisha 760010, India

### Supplementary data

**Figure S1**

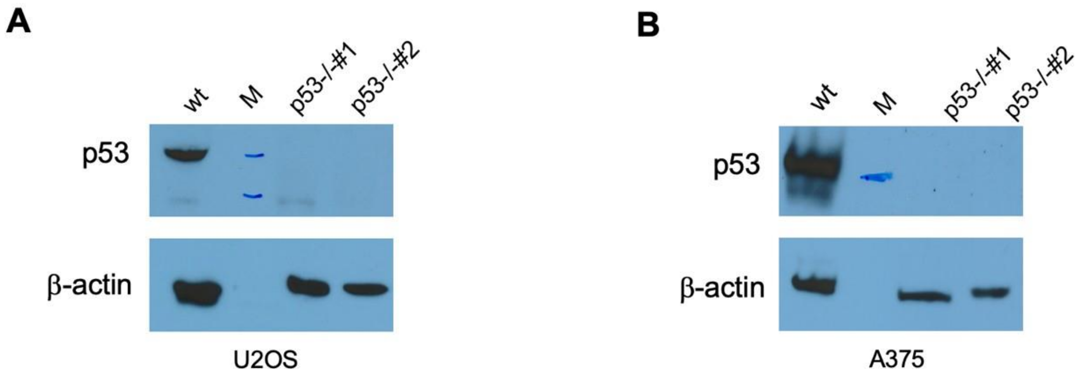

**Figure S1: p53$^{-/-}$ (ko) validation.**

(A-B) In U2OS and A375 cell lines, p53 was knocked out using CRISPR-Cas9. Western blot of U2OS and A375 cells lines for p53 left panel (A) using monoclonal p53 antibody and right panel (B) as β-actin is presented as a loading control. The two individual p53 knock out clones were validated in both cell lines. Ko#1 was used throughout the studies in both the cell lines. wt=wild type, M=marker

#### Figure S2

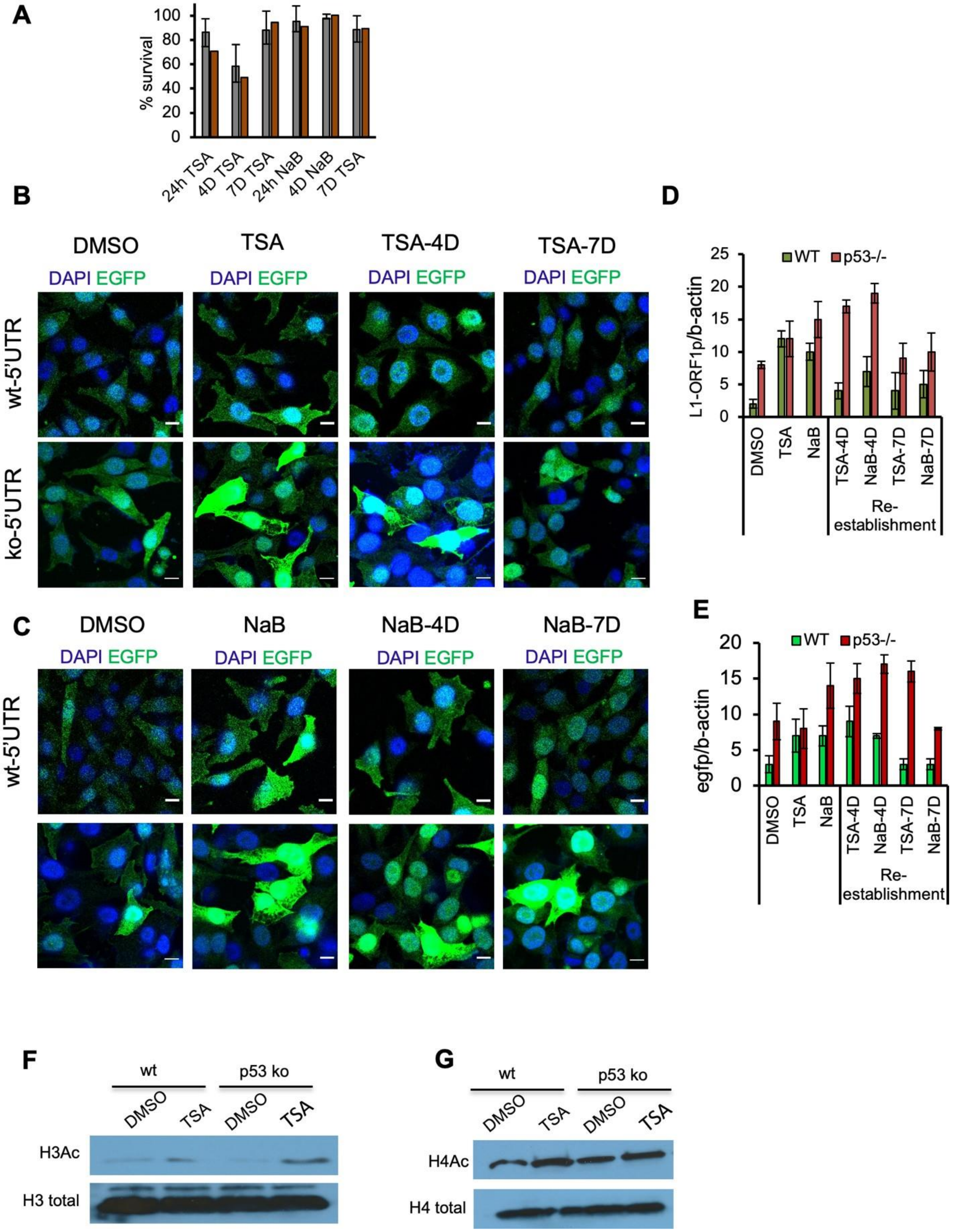
A
wt-5'UTR
p53-/-5'UTR
% survival
24h TSA
4D TSA
7D TSA
24h NaB
4D NaB
7D TSA
B
DMSO
TSA
TSA-4D
TSA-7D
DAPI EGFP
wt-5'UTR
ko-5'UTR
C
DMSO
NaB
NaB-4D
NaB-7D
DAPI EGFP
wt-5'UTR
D
WT
p53-/-
L1-ORF1p/b-actin
DMSO
TSA
NaB
TSA-4D
NaB-4D
TSA-7D
NaB-7D
Re-establishment
E
WT
p53-/-
egfp/b-actin
Re-establishment
F
wt
p53 ko
DMSO
TSA
H3Ac
H3 total
G
wt
p53 ko
DMSO
TSA
H4Ac
H4 total

20

**Figure S2: p53 mediates silencing of L1 in wt in response to HDACi treatment.**

(A) The percentage survival of A375-L1-5'UTR-eGFP reporter wt (wt-5'UTR) and p53$^{-/-}$ (p53$^{-/-}$ -5'UTR) cells upon HDACi drug treatment at 24 h of drug treatment and followed by drug withdrawal of 3 and 6 days. Measurement of luminescent signals was conducted at 1 day, 4 days and 7 days post HDACi drug treatment. Data are shown as the percentage of cell viability compared with the DMSO control, which was defined as 100%. Bar graph displayed the mean from three Biological replicates each with 3 technical replicates.

(B) Refer to **Figure 4A** for the schematics of HDACi treatment and withdrawal. Confocal imaging of GFP was conducted in stable cell lines of wt and p53$^{-/-}$ A375 carrying L1-5'UTR-eGFP reporter, treated with either DMSO, TSA, or NaB for 24 h, followed by withdrawal for 3 and 6 days. Samples were collected 1 day after HDACi exposure, and at 4 and 7 days for confocal imaging. Representative of three independent immunofluorescence experiments is shown here. Activation of GFP was observed upon TSA treatment, followed by subsequent re-silencing of GFP in wt cells after the withdrawal experiment.

(C-D) RT-qPCR of L1-ORF1 (E) and (F) GFP transcripts in wt and p53$^{-/-}$ L1-5'UTR cells in DMSO (vehicle) or TSA/NaB treated for 24 h and drug withdrawal after 3 and 6-days. Samples were collected for analyses at 1 day, 4 days and 7 days. Shown is the mean± S.D. from three independent biological replicates (n=3). The primers for qPCR analysis of ORF1 and egfp transcripts are shown in (**Supplemental Table 1**)

(E-F) Western blotting of H3Ac and total H3 levels (E) and H4Ac and H4 total levels (F) in A375 wt and p53$^{-/-}$ cell lines carrying with L1-5'UTR-egfp reporter upon treatment either with DMSO or TSA for 24 h.

**Figure S3**

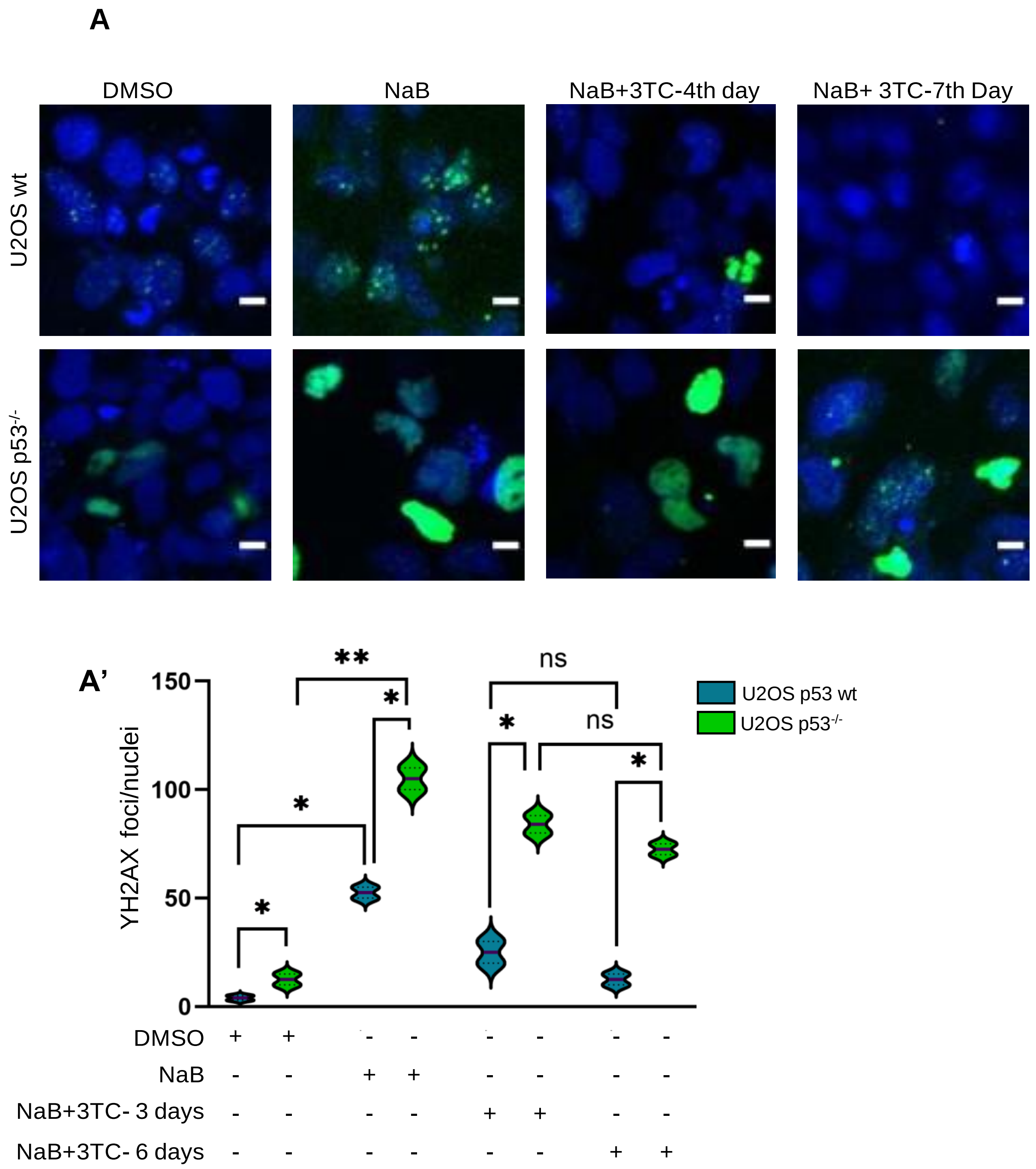

**Figure S3: Effect of 3TC on HDACi activated U2OS wt and p53$^{-/-}$**

(A) Confocal imaging of yH2AX foci (green) in wt, and p53$^{-/-}$ U2OS treated with either DMSO or TSA for 24 h followed by 3TC treatment of 3 and 6 days Samples were imaged at 1 Day (24 h treated), 4 days and 7 days of post drug exposure . Shown here is the representative confocal imaging of 3 independent immunofluorescence imaging.  Scale bar denotes 20um. Blue denotes nucleus.

(A’) The Violin plot shows the quantified yH2AX foci upon TSA treatment followed and 3TC treatment of 3 and 6 days. The bold dotted line represents the mean foci from 50 different nuclei. The quantification has been done from 3 different biological replicates. *p<0.05, **p<0.01, ns=non-significant (Mann-Whitney U test).

**Figure S4**

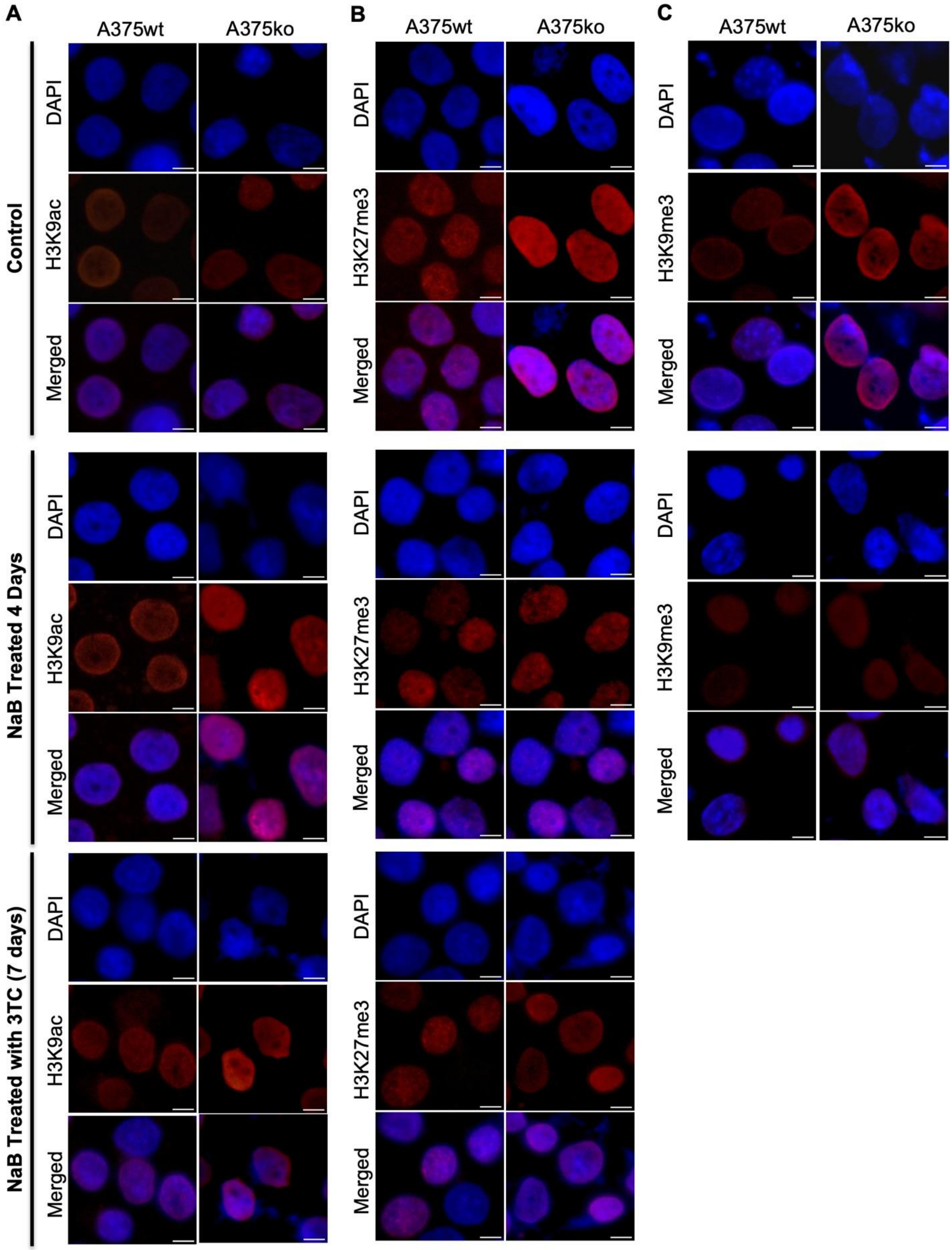
A
A375wt
A375ko
Control
NaB Treated 4 Days
NaB Treated with 3TC (7 days)
DAPI
H3K9ac
Merged
B
A375wt
A375ko
DAPI
H3K27me3
Merged
C
A375wt
A375ko
DAPI
H3K9me3
Merged

57

**Figures S4: Sodium Butyrate-mediated HDAC inhibition increases global histone acetylation with loss of histone repressive marks**

(A-C) Represented Immunofluorescence images show epigenetic marks in control (DMSO) and NaB treated samples followed after drug withdrawal. The marks include (A) H3K9Ac (activation mark) (B) H3K27Me3 (repressive mark) and (C) H3K9Me3 (repressive mark). The data represents comprehensive data from 3 biological replicates. Scale bar indicate 50uM.

**Figure S5**

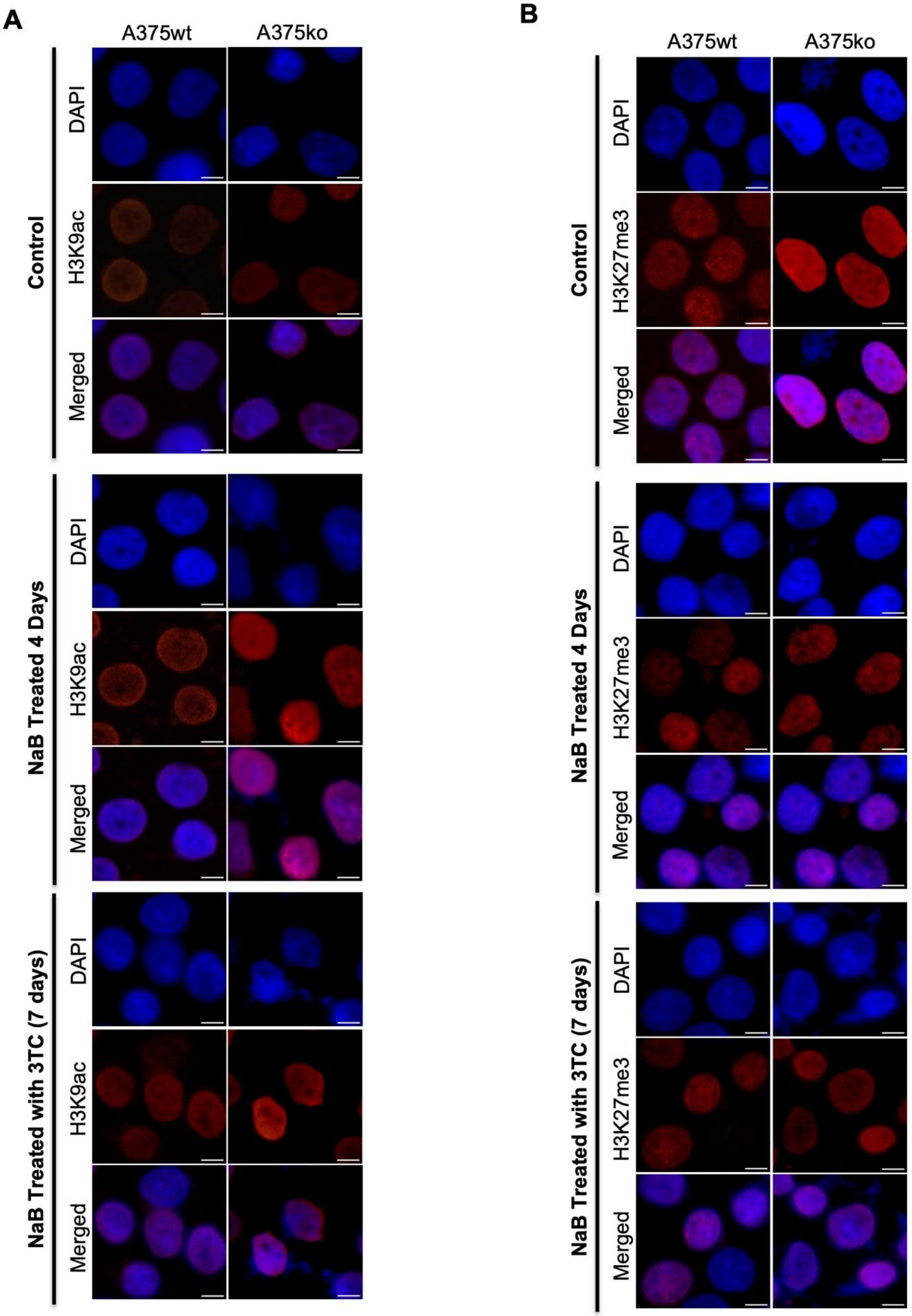
A
A375wt
A375ko
Control
NaB Treated 4 Days
NaB Treated with 3TC (7 days)
DAPI
H3K9ac
Merged
B
A375wt
A375ko
Control
NaB Treated 4 Days
NaB Treated with 3TC (7 days)
DAPI
H3K27me3
Merged

64

**Figures S5: Effect of 3TC on  global on epigenetic histone marks**

(A-C) Represented Immunofluorescence images show epigenetic marks in control (DMSO) and NaB treated samples followed by 3TC treatment. The marks include (A) H3K9Ac (activation mark) (B) H3K27Me3 (repressive mark). The data represents comprehensive data from 3 biological replicates. Scale bar indicate 50uM.

**Supplementary Table 1**

| | Primer name | Primer sequence |
|---|---|---|
| **sgRNA** | p53 sgRNA | GCTTGTAGATGGCCATGGCG |
| | huAAVS1 sgRNA | GTTAATGTGGCTCTGGTTCT |
| **AAVS1 Cloning** | L1-5’UTR-FseI Forward | ATATAATTGGCCGGCCGGGGGAGGAGCCAAG ATG GCCGAATA |
| | L1-5’UTR-AfeI Reverse | CCAACCAGCGCTCTTTGTGGTTTTATCTACTTT TGGTCTTTGATGATGGTGATGTACAGA |
| **qRT-PCR (real time)** | b-actin Forward | CCGCGAGAAGATGACCCAGAT |
| | b-actin Reverse | CGTTGGCACAGCCTGGATAGCAACG |
| | eGFP Forward | GACTTCAAGGAGGACGGCAA |
| | eGFP Reverse | TCTCGTTGGGGTCTTTGCTC |
| | 5'UTR PP1 Forward | ACAGTGGGGCAAGCTGG |
| | 5'UTR PP1 Reverse | TCACCCCTTTCTTTGACTCG |
| | L1-ORF1 Forward | GGTTACCCTCAAAGGAAAGCC |
| | L1-ORF1 Reverse | ATGAAGCTTAGTTTGGCTGG |
| | L1-ORF2 Forward | AAATGGTGCTGGGAAAACTG |
| | L1-ORF2 Reverse | GTTTTGGACATGAAGTCCTTGC |
| | sting Forward | CCTGAGTCTCAGAACAACTGCC |
| | sting Reverse | GGTCTTCAAGCTGCCCACAGTA |
| | rb Forward | TGATGAGTGGGTGAAGGCTGAC |
| | rb Reverse | CCCGTTTTGACTTCTGCCTCTC |
| | rad50 Forward | CCAGACCCAGCTCCTTTACC |
| | rad50 Reverse | CACTGCGACACCAAACTCATC |
| | mcp2 Forward | TATCCAGAGGCTGGAGAGCTAC |
| | mcp2 Reverse | TGGAATCCCTGACCCATCTCTC |
| | rpl13A (positive R-loop locus) Forward | CTCCTCCTGCCTGCATTCTT |
| | rpl13A (positive R-loop locus) Reverse | CACACTTCCTGCCCACTTGT |
| | znf544 (negative R-loop locus) Forward | ATGACTGGTCTTCCCGCTTG |
| | znf544 (negative R-loop locus) Reverse | CGCTTCTGAACCTGACACCA |
| | Brca1 Forward | CTGAAGACTGCTCAGGGCTATC |
| | Brca1 Reverse | AGGGTAGCTGTTAGAAGGCTGG |
| | Il-6 Forward | AGACAGCCACTCACCTCTTCAG |
| | Il-6 Reverse | TTCTGCCAGTGCCTCTTTGCTG |

| | | |
|---|---|---|
| **L1 ORF1,EN, RT Cloning** | L1-ORF1p Forward (EcoRI) | TTCGAATTCATGGGGAAAAAACAGAAC |
| | L1-ORF1p Reverse (BamHI) | GATGGATCCCATTTTGGCATGATTTTG |
| | L1 Endonuclease Forward (EcoRI) | TTCGAATTCATGACAGGATCAAATTCA |
| | L1 Endonuclease Reverse (KpnI) | CGCGGTACCAATCCTGAGTTCTAGTTT |
| | L1 Reverse transcriptase Forward (EcoRI) | CTTCGAATTCATGTTCAATAGAAAAAGAG |
| | L1 Reverse transcriptase Reverse (KpnI) | CGCGGTACCAAGTTGGATTCCTAGGTA |

### Supplementary Table 2

| | LINE1-ORF1P STELLARIS PROBE SEQUENCES |
|---|---|
| 1 | CTGTTCTGTTTTTTCCCCAT |
| 2 | CGTTTTAGAGTTTCCAGTTT |
| 3 | TTGGAGGAGGAGAGGCGCTC |
| 4 | GCTGGTGAGGAACTGCGTTC |
| 5 | CTCCATCCAGCTTTGTTCTG |
| 6 | CTCTCAGCTCATCAAAATCA |
| 7 | TTTGATCGTCTGAAGCCTTC |
| 8 | GTCCTCCCGTAGCTCAGAGT |
| 9 | CTTCTTTGCCTTTGGTTTGA |
| 10 | AAATTTTTTTCAAAGTTTTC |
| 11 | TATTCTAGTTATACATTCTT |
| 12 | TTAAGCACTTCTCTGTATTG |
| 13 | GTTTTCAGCTCCATCAGCTC |
| 14 | TTCACGTAGTTCTCGAGCCT |
| 15 | GGCTCCTGAGGCTTCTGCAT |
| 16 | CTTTCTTCCAGTTGATCGCA |
| 17 | TTCATCTTCCATTGCTGATA |
| 18 | CTTCTCGCTTCATTTCATTC |
| 19 | ATTCTTTTTTCTCTAAACTT |
| 20 | GAGGCTTTGCTCATTTCTTT |
| 21 | TCACATAGTCCCATATTTCT |
| 22 | ATCAGACGTAGATTTGGTCT |
| 23 | CACATCACTTTCAGGTACAC |
| 24 | TTTCCAACTTGGTTCCATTC |
| 25 | TGGATAATATCCTGCAGAGT |
| 26 | TGCTAGATTGGGGAAGTTCT |
| 27 | TGAATCTGAACGTTGGCCTG |
| 28 | TTGTGGCGTTCTCTGTATTT |
| 29 | TCGCTCTTCTCGAGGAGTAT |
| 30 | CTGACAATTATGTGTCTTGG |

| | |
|---|---|
| 31 | CTTCATTTCAACTTTGGTGA |
| 32 | TGGCTGCCCTTAACATTTTT |
| 33 | AGGGTAACCCGACCTTTCTC |
| 34 | TAGTCTGATGGGCTTTCCTT |
| 35 | TTTCTGCCGAGAGATCCGCT |
| 36 | CACTCCCTTCTGGCTTGTAG |
| 37 | AAGAATGTTGAATATTGGCC |
| 38 | TGGGTTGAAAATTCTTTTCT |
| 39 | CTTAGTTTGGCTGGATATGA |
| 40 | TATTTCTCCTTCACTTATGA |
| 41 | CATTTGCTTGTCTATAAAGT |
| 42 | TGGTGGTGACAAAATCTCTC |
| 43 | AGGAGCTCTTTTAGGGCAGG |
| 44 | TTCCATGTTTAGCGCTTCCT |
| 45 | GCGGCTGGTACCGGTTGTTC |
| 46 | TACATTTTGGCATGGTTTTG |

| | **Gene Sequences** | |
|---|---|---|
| 1 | ORF1p | ATGGGGAAAAAACAGAACAGAAAAACTGGAAACTCTAAAACGCAGA<br>GCGCCTCTCCTCCTCCAAAGGAACGCAGTTCCTCACCAGCAACAGA<br>ACAAAGCTGGATGGAGAATGATTTTGACGAGCTGAGAGAAGAAGGC<br>TTCAGACGATCAAATTACTCTGAGCTACGGGAGGACATTCAAACCAA<br>AGGCAAAGAAGTTGAAAACTTTGAAAAAAATTTAGAAGAATGTATAA<br>CTAGAATATCCAATACAGAGAAGTGCTTAAAGGAGCTGATGGAGCT<br>GAAAACCAAGGCTCGAGAACTACGTGAAGAATGCAGAAGCCTCAGG<br>AGCCGATGCGATCAACTGGAAGAAAGGGTATCAGCAATGGAAGATG<br>AAATGAATGAAATGAAGCGAGAAGGGAAGTTTAGAGAAAAAAGAATA<br>AAAAGAAATGAGCAAAGCCTCCAAGAAATATGGGACTATGTGAAAAG<br>GCCAAATCTACGTCTGATTGGTGTACCTGAAAGTGATGTGGAGAAT<br>GGAACCAAGTTGGAAAACACTCTGCAGGATATTATCCAGGAGAACTT<br>CCCCAATCTAGCAAGGCAGGCCAACGTTCAGATTCAGGAAATACAG<br>AGAACGCCACAAAGATACTCCTCGAGAAGAGCAACTCCAAGACACA<br>TAATTGTCAGATTCACCAAAGTTGAAATGAAGGAAAAAATGTTAAGG<br>GCAGCCAGAGAGAAAGGTCGGGTTACCCTCAAAGGGAAGCCCATC<br>AGACTAACAGCGGATCTCTCGGCAGAAACCCTACAAGCCAGAAGAG<br>AGTGGGGGCCAATATTCAACATTCTTAAAGAAAAGAATTTTCAACCC<br>AGAATTTCATATCCAGCCAAACTAAGCTTCATAAGTGAAGGAGAAAT<br>AAAATCCTTTACAGACAGGCAAATGCTGAGAGATTTTGTCACCACCA<br>GGCCTGCCCTAAAAGAGCTCCTGAAGGAAGCGCTAAACATGGAAAG<br>GAACAACCGGTACCAGCCGCTGCAAAATCATGCCAAAATG |

| | | |
|---|---|---|
| 2 | EN domain | ATGACAGGATCAAATTCACACATAACAATATTAACTTTAAATGTAAATGGACTAAATTCTCCAATTAAAAGACACAGACTGGCAAGTTGGATAAAGAGTCAAGACCCATCAGTGTGCTGTATTCAGGAAACCCATCTCACGTGCAGAGACACACATAGGCTCAAAATAAAAGGATGGAGGAAGATCTACCAAGCAAATGGAAAACAAAAAAAGGCAGGGGTTGCAATCCTAGTCTCTGATAAAACAGACTTTAAACCAACAAAGATCAAAAGAGACAAAGAAGGCCATTACATAATGGTAAAGGGATCAATTCAACAAGAGGAGCTAACTATCCTAAATATTTATGCACCCAATACAGGAGCACCCAGATTCATAAAGCAAGTCCTGAGTGACCTACAAAGAGACTTAGACTCCCACACATTAATAATGGGAGACTTTAACACCCCACTGTCAATATTAGACAGATCAACGAGACAGAAAGTCAACAAGGATACCCAGGAATTGAACTCAGCTCTGCACCAAACAGACCTAATAGACATCTACAGAACTCTCCACCCCAAATCAACAGAATATACATTTTTTTCAGCACCACACCACACCTATTCCAAAATTGACCACATAGTTGGAAGTAAAGCTCTCCTCAGCAAATGTAAAAGAACAGAAATTATAACAAACTATCTCTCAGACCACAGTGCAATCAAACTAGAACTCAGGATT |
| 3 | RT domain | TCAATAGAAAAAGAGGGAATCCTCCCTAACTCATTTTATGAGGCCAGCATCATTCTGATACCAAAGCCGGGCAGAGACACAACCAAAAAAGAGAATTTTAGACCAATATCCTTGATGAACATTGATGCAAAAATCCTCAATAAAATACTGGCAAACCGAATCCAGCAGCACATCAAAAAGCTTATCCACCATGATCAAGTGGGCTTCATCCCTGGGATGCAAGGCTGGTTCAATATACGCAAATCAATAAATGTAATCCAGCATATAAACAGAGCCAAAGACAAAAACCACGTGATTATCTCAATAGATGCAGAAAAAGCCTTTGACAAAATTCAACAACCCTTCATGCTAAAAACTCTCAATAAATTAGGTATTGATGGGATGTATCTCAAAATAATAAGAGCTATCTATGACAAACCCACAGCCAATATCATACTGAATGGGCAAAAACTGGAAGCATTCCCTTTGAAAACTGGCACAAGACAGGGATGCCCTCTCTCACCACTCCTATTCAACATAGTGTTGGAAGTTCTGGCCAGGGCAATCAGGCAGGAGAAGGAAATAAAGGGTATTCAATTAGGAAAAGAGGAAGTCAAATTGTCCCTGTTTGCAGACGACATGATTGTTTATCTAGAAAACCCCATCGTCTCAGCCCAAAATCTCCTTAAGCTGATAAGCAACTTCAGCAAAGTCTCAGGATACAAAATCAATGTACAAAAATCACAAGCATTCTTATACAACAACAACAGACAAACAGAGAGCCAAATCATGGGTGAACTCCCATTCACAATTGCTTCAAAGAGAATAAAATACCTAGGAATCCAACTT |
